# Population genomics of the endangered scaly-foot snail defines conservation units amid deep-sea mining threats in Indian Ocean vents

**DOI:** 10.1101/2025.08.02.668274

**Authors:** Xi Chen, Siyeong Jun, Chong Chen, Ran Liu, Ting Xu, Hongyin Zhang, Yue Gao, Xing He, Xu Liu, Kexin Gao, Xinyu Gu, Heeseung Yeum, Dongyoung Kim, Choongwon Jeong, Yadong Zhou, Ken Takai, Yan Wang, Pei-Yuan Qian, Yong-Jin Won, Jin Sun

**Affiliations:** Key Laboratory of Evolution & Marine Biodiversity (Ministry of Education) and Institute of Evolution & Marine Biodiversity, Ocean University of China, Qingdao 266003, China; Laboratory for Marine Biology and Biotechnology, Qingdao Marine Science and Technology Centre, Laoshan Laboratory, Qingdao 266237, China; Department of Life Science, Division of EcoScience, Ewha Womans University, Seoul 03760, South Korea; X-STAR, Japan Agency for Marine-Earth Science and Technology (JAMSTEC), 2-15 Natsushima-cho, Yokosuka, Kanagawa 237-0061, Japan; Southern Marine Science and Engineering Guangdong Laboratory (Guangzhou), Guangzhou, China; Department of Ocean Science, The Hong Kong University of Science and Technology, Hong Kong, China; State Key Laboratory of Submarine Geoscience, Second Institute of Oceanography, Ministry of Natural Resources, Hangzhou, China; Key Laboratory of Marine Ecosystem Dynamics, Second Institute of Oceanography, Ministry of Natural Resources, Hangzhou, China; School of Biological Sciences, Seoul National University, Seoul 08826, South Korea; Institute for Data Innovation in Science, Seoul National University, Seoul 08826, South Korea

## Abstract

Hydrothermal vents in the Indian Ocean face imminent deep-sea mining, yet lack baseline high-resolution connectivity data for their endemic fauna. We present the first population genomic study of the endangered scaly-foot snail (*Chrysomallon squamiferum*), an iconic species distributed across three vent biogeographic provinces. Analysing 125 individuals from eight vent fields using 14 million single-nucleotide polymorphisms, we identify five genetic groups that warrant recognition as evolutionarily significant units. Demographic modelling reveals critical contributions from extinct or unsampled ‘phantom populations’ to the contemporary genetic structure. Together with physical ocean modelling, we show that the deep currents drive asymmetric south-to-north gene flows, while transform faults act as dispersal barriers. We propose Longqi-Duanqiao fields on the Southwest Indian Ridge and Wocan field on the Carlsberg Ridge as two isolated populations prioritised for protection. Given its exceptional adaptations, wide distribution, and public recognition, the scaly-foot snail is well-positioned to serve as both a flagship and umbrella species to ensure the survival of a broader suite of vent biodiversity. As the International Seabed Authority finalises its Mining Code in 2025, our findings provide essential genomic evidence to mitigate the impacts of mining on genetic diversity and inform transboundary conservation strategies for this vulnerable ecosystem.

## Introduction

The seminal theory of island biogeography by MacArthur and Wilson posits that geographic isolation, mediated by uninhabitable matrices, drives diversification and speciation in insular populations – ultimately culminating to forming the global biodiversity at population to ecosystem scales^1^. Analogous “island-like” dynamics operate in specialised habitats of the vast, seemingly interconnected deep ocean, where systems like hydrothermal vents function as biological islands for an endemic fauna that cannot survive elsewhere. Since their landmark discovery in 1977, vents have captivated scientists as biological hot spots with a high biomass of unique organisms, such as the giant tubeworm *Riftia pachyptila* and Pompeii worm *Alvinella pompejana*, the most heat-tolerant animal^2^. These ‘extreme’ ecosystems are defined by high concentrations of reducing chemicals and heavy metals, dramatic second-scale fluctuations in environmental parameters, and immense hydrostatic pressure^3^. The secret to their success? Bacterial primary production in the form of chemosynthesis using reducing chemicals, often manifesting as close-knit relationships between animals and symbionts.

So far, over 650 active hydrothermal vents have been discovered worldwide^4^, mostly in mid-ocean ridges, volcanic arcs, and back-arc basins; vents from each geographic locality differ in faunal composition, partitioning them into distinct biogeographic provinces^5, 6^. Vents usually function as metapopulations – networks of spatially distinct subpopulations interconnected by episodic larval exchange – as most vent species have a planktonic larval stage^7^. Population genetics and genomics are powerful proxies for deciphering how vent-endemic species maintain their distribution ranges through both historical and contemporary effective larval dispersal^8, 9, 10, 11^. These approaches illuminate population connectivity and potential speciation processes by revealing genomic divergence across discontinuous habitats, and are important since the direct quantification by tracking microscopic larvae is practically impossible in the vast ocean.

The dispersal dynamics of vent-endemic species are governed by an interplay between intrinsic biological traits and external physical drivers. A species’ inherent dispersal potential is first set by its life-history, such as larval type, pelagic larval duration (PLD), and physiological tolerance. Along mid-ocean ridges, buoyancy-driven currents aligned with ridge axes often serve as dispersal “highways” that enable long-distance connectivity between vent fields along the ridge^12^, while in back-arc basins a largely enclosed circulation tends to retain larvae within local basins, promoting genetic isolation^13^. Regional heterogeneity further arises from other drivers such as ridge spreading rates, which modulate vent field density as well as longevity, and geofluid chemistry differences leading to the colonisation of species with different physiological tolerances and ecological niches^14, 15, 16^. Together, these factors interact with species-specific larval ecology to generate complex biogeographic mosaics where isolation-by-distance frequently coexists with abrupt genetic discontinuities across dispersal barriers formed by transform faults or oceanographic fronts^8^. Understanding the role and contribution of each driver is essential for reconstructing population histories and predicting vulnerability to both natural and anthropogenic disturbances^17^.

Hydrothermal vents in the Indian Ocean are imminently threatened by deep-sea mining, with four nations — China, Germany, India, and South Korea — together holding five exploratory licenses from the International Seabed Authority (ISA) that target almost all confirmed active vent systems within the Indian Ocean [except one within the Mauritius Exclusive Economic Zone (EEZ)]^18^. As the ISA moves towards finalising its Mining Code before 2026, the need for scientific data to inform conservation and management strategies has become critically urgent to mitigate potential biodiversity loss^19, 20^. At least 15 confirmed hydrothermal vent fields exist on Indian Ocean mid-ocean ridges^21^, partitioned into three biogeographic provinces: the Carlsberg Ridge (CR) province, the Central Indian Ridge - northern Southwest Indian Ridge (CIR-nSWIR) province (including all CIR vents plus Tiancheng vent field on the nSWIR), and the southern Southwest Indian Ridge (sSWIR) province (including Longqi and Duanqiao fields)^22^. These ridges differ in spreading rate leading to divergent landscapes: the ultra-slow spreading SWIR fosters widely spaced, long-lived vents with high isolation potential, while faster-spreading CIR and CR support denser vent fields with theoretically greater connectivity^23^. With India applying for ISA exploratory license on the CR, all three vent provinces are now eyed for mining^24^. Compared Pacific and Atlantic counterparts, the baseline biodiversity and dispersal data for Indian Ocean vents are lacking due to remoteness and historical lack of research focus^25, 26^. Critically, no population genomic studies exist for any Indian Ocean vent species – leading to a restricted knowledge of faunal connectivity that limits our capacity to inform conservation strategies. This significant knowledge gap in how genetic diversity is generated, maintained, and segregated across these geodynamically distinct vents needs to be filled in order to decipher large-scale biogeographic patterns in the deep Indian Ocean and also urgently needed for management^27^.

The scaly-foot snail *Chrysomallon squamiferum* is an iconic species endemic to Indian Ocean vents, famed for its ‘ironclad’ scale-armour that has inspired advanced protective materials engineering^28, 29, 30^ and the production of iron sulfide nanoparticles^31^ – although the scales actually function as sites of sulfur detoxification^31^. In 2019, this species became the first hydrothermal vent species assessed as Endangered from upcoming deep-sea mining in the IUCN (International Union for Conservation of Nature) Red List^32^ and is the prime candidate for a flagship and umbrella species in conservation actions. This peltospirid snail depends entirely on endosymbiotic gammaproteobacteria for nutrition via sulfur oxidation^33, 34^, making it completely reliant on the vent habitat. With lecithotrophic (non-feeding, yolk-dependent) larvae, it is thought to disperse via deep-ocean currents^35^ and is present across all three Indian Ocean biogeographic provinces from sSWIR to CR making it an ideal model for ridge-scale connectivity studies to inform conservation policy through the identification of evolutionarily significant units^35, 36^. However, only limited population-level studies have been done using the barcoding fragment of the partial mitochondrial COI (cytochrome oxidase *c* subunit I) gene^22, 35, 37^, which indicated population structuring across ridges. The limited resolution of this single marker made it impossible to delineate complex, fine-scale genetic structure or reliable quantification of gene flow and demographic dynamics.

Here, we present the first comprehensive population genomic analysis of vent fauna in the Indian Ocean, using 125 individuals of *C. squamiferum* sampled across eight vent fields spanning all three Indian Ocean biogeographic provinces and all five ISA contract areas (Figure 1a). By coupling genome-wide SNP markers with physical oceanographic modelling, we aim to: (1) quantify basic genetic parameters and identify potential dispersal barriers structuring populations; (2) reconstruct contemporary and ancient gene flow patterns to elucidate the historical processes shaping Indian Ocean vent biogeography; (3) characterise asymmetric migration and demographic connectivity to identify evolutionarily significant units and source-sink dynamics critical for conservation planning; and (4) identify vent fields that warrant prioritised protection.

**Figure 1.**
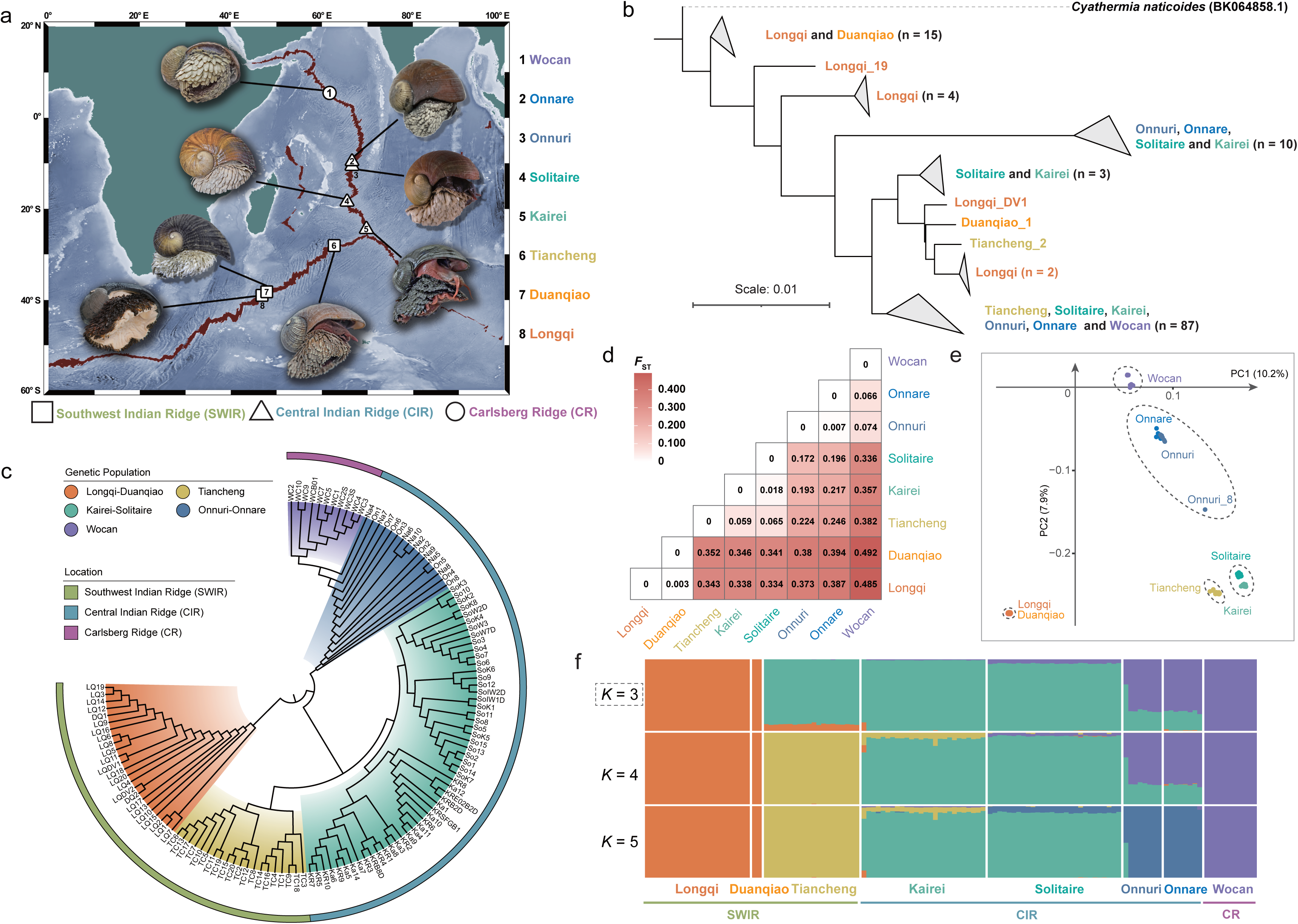
Population structure and genetic divergence of the scaly-foot snail, *Chrysomallon squamiferum*. **a.** Geographic distribution of the vent fields where the scaly-foot snails were collected, and photographs of a representative specimen from each; the eight hydrothermal vent areas are distributed along the Southwest Indian Ridge, Central Indian Ridge, and Carlsberg Ridge. **b.** Phylogenetic tree constructed using the Maximum Likelihood (ML) method based on all 13 mitochondrially encoded genes; the vent snail *Cyathermia naticoides* from Neomphalidae, the sister-family of Peltospiridae containing *C. squamiferum*, served as the outgroup. **c.** Phylogenetic tree of scaly-foot snail individuals constructed using the distance-based method with SNP data. **d.** Pairwise Hudson’s *F*_ST_ values between local population pairs, indicating population genetic differentiation estimated using PLINK2. *F*_ST_ values range from 0 to 1; values closer to 0 indicate a lower degree of genetic differentiation between populations. **e.** Genetic discrepancies among scaly-footsnail populations revealed by principal component analysis (PCA). The results show low genetic differentiation between the Longqi and Duanqiao populations, the Solitaire and Kairei populations, and the Onnuri and Onnare populations. These pairs were combined into genetic groups, indicated by dashed circles. **f.** Population structure and individual ancestry detected using ADMIXTURE, assuming three to five predefined genetic groups. The optimal number of groups (*K* = 3) is indicated by a dotted box.

## Results

### Sample collection and detection of genome-wide SNPs

A total of 125 scaly-foot snails were collected from eight hydrothermal vent fields in the Indian Ocean between 2013 and 2023 (Figure 1a; Table 1), and we generated whole genome resequencing data (average depth: 38.72*×*, range: 18.25–212.70*×*, Supplementary Data 1) for them. Our sampling localities cover three mid-ocean ridges of the Indian Ocean and the entire known range of the species^22^, including Longqi, Duanqiao, and Tiancheng vent fields on the SWIR; Kairei, Solitaire, Onnuri, and Onnare vent fields on the CIR; and the Wocan vent field on the CR. The sampling sites spanned 6387 kilometres along the ridge axis and between 1780 and 2993 metres in depth. Two distinct pipelines (i.e., GATK and bcftools) were used to detect genome-wide single-nucleotide polymorphisms (SNPs). After strict quality control, the GATK pipeline produced 18,939,600 SNPs, while the bcftools pipeline yielded 24,309,642 SNPs. A total of 14,309,443 high-confidence SNPs identified by both pipelines, including 10,338,002 rare SNPs (MAF<0.05) and 3,971,441 conventional SNPs were used for downstream analyses.

**Table 1.**
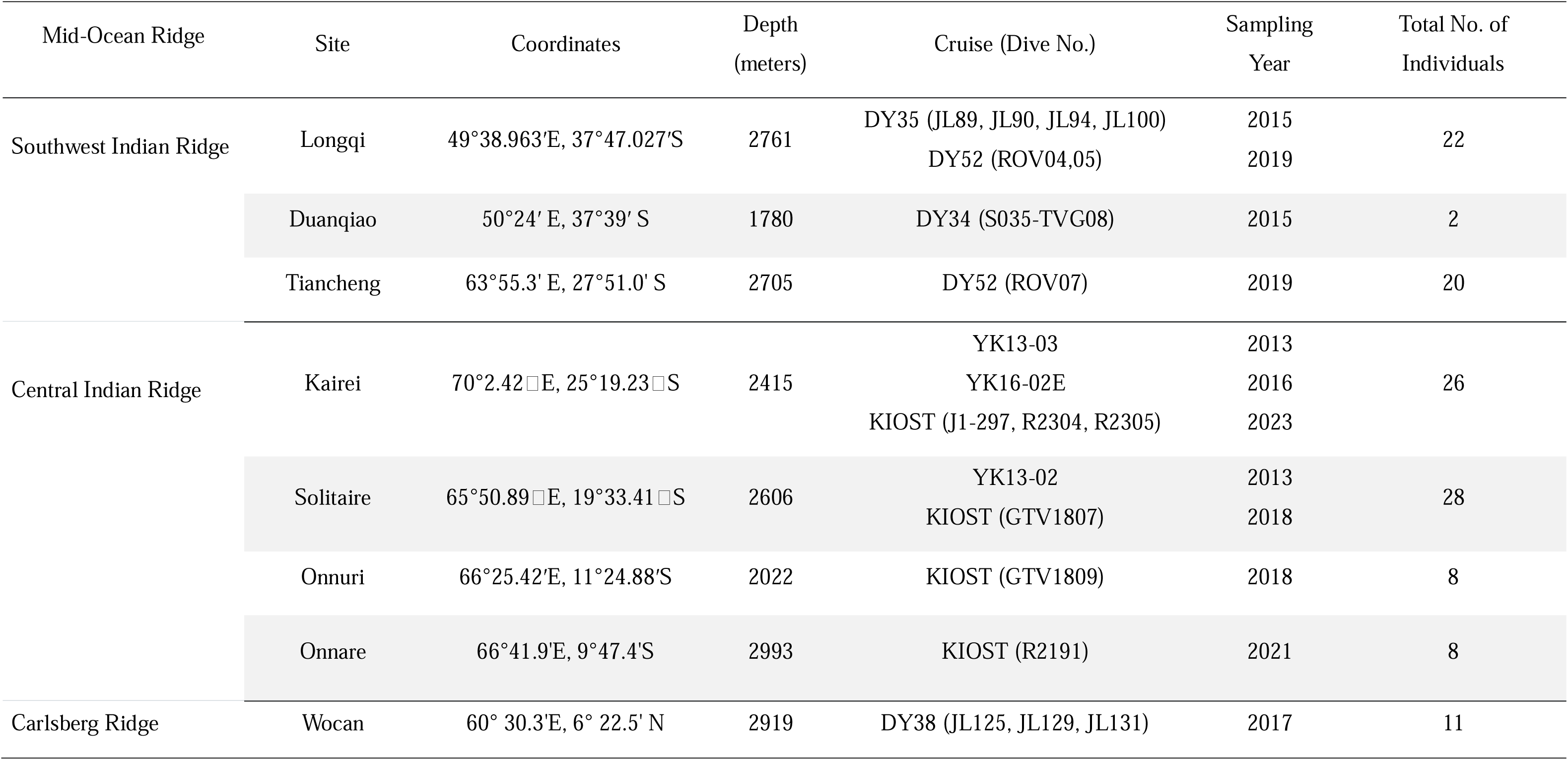
Sampling data of scaly-foot snails used in this study.

### Population structure

To decipher the hierarchical population genetic structure, we computed the pairwise *F*_ST_ (Figure 1d) inbreeding coefficients (*F*) (Supplementary Data 2) and performed principal component analyses (PCA) (Figure 1e) using the conventional SNPs. Pairwise *F*_ST_ values between all population pairs ranged between 0.003 to 0.492 (Figure 1d, Supplementary Data 3). Notably, Longqi and Duanqiao showed consistently high genetic differentiation from the rest of the populations (*F*_ST_ range: 0.343–0.492). In contrast, the three population pairs with the lowest differentiation were Longqi and Duanqiao, Kairei, and Solitaire, as well as Onnuri and Onnare (*F*_ST_ range: 0.003–0.018). Generally, a significant positive correlation between *F*_ST_ and geographical distance along the ridges, indicative of an isolation-by-distance pattern, was observed (*R^2^* = 0.93, *P*-value = 7.2 **×** 10^-10^) (Figure S1), except Onnuri, Onnare and Wocan, despite being geographically more distant from each other than from Solitaire, exhibited lower genetic differentiation (*F*_ST_ = 0.066–0.074 vs. 0.172–0.196). The scaly-foot snails of Longqi and Duanqiao showed the highest and significantly elevated inbreeding coefficients (*F* = 0.37–0.44 vs. 0.04–0.36), a finding that aligns with their strong genetic differentiation from other populations as measured by *F*_ST_.

In the PCA plot, individuals clustered by vent field, in a pattern generally consistent with their geographical locations. According to the first two principal components, the snails formed five major genetic groups, including Longqi-Duanqiao, Tiancheng, Kairei-Solitaire, Onnuri-Onnare, and Wocan. One individual from Onnuri (On8) showed considerable genetic divergence from the other Onnuri individuals, positioning it between Onnuri and Kairei-Solitaire. This hybrid-like individual was consistently identified in subsequent Admixture analyses, irrespective of *K* numbers (Figure 1f). In our Admixture analyses with the number of genetic groups (*K*) set between 2 to 8 (Figure 1f, Figure S2a), *K* = 3 was identified as the optimal (Figure S2b) which showed likely introgression between Longqi-Duanqiao and Tiancheng, as well as between Onnuri-Onnare and Kairei-Solitaire. At *K* = 4, the introgression from Tiancheng was observed for Kairei. At *K* = 5, the eight populations of scaly-foot snails were divided into five genetic groups, consistent with the results of PCA (Figure 1d). Taking these evidences together, hereafter we treat the Longqi-Duanqiao pair as a single genetic group, as well as the Kairei-Solitaire pair and the Onnuri-Onnare pair.

### Relationships among groups

A phylogenetic tree constructed based on 13 protein-coding genes of our scaly-foot snail mitogenomes revealed that the Longqi population formed the earliest-diverging lineage, sister to all remaining populations (Figure 1b and Figure S3). Results from our badMIXTURE analysis supported this, as the Longqi-Duanqiao populations exhibited the lowest residuals after fitting optimal ancestral palettes (Figure S4). This suggests they encompass more ancestry loci of all contemporary populations, implying they share close kinship with the ancestral population. Furthermore, these results also indicate the Longqi-Duanqiao population formed without ‘phantom’ populations (either now-extinct or extant but unsampled) or going through severe bottlenecks (Figure S4). Population history modelling consistently supported this relationship: qpGraph simulations rotating different genetic groups as outgroups revealed that models positioning Longqi-Duanqiao as the outgroup yielded significantly lower scores compared to observed data across 0 to 4 assumed introgressions (the lower the score, the better the fit between model and data, *P* <0.01, Figure S5a, S5b). Similarly, Fastsimcoal2 simulations confirmed Longqi-Duanqiao as the optimal outgroup (Δlikelihood score = 2,165,338, *P* <0.01, Figure S6). Collectively, these results overwhelmingly support the Longqi-Duanqiao genetic group as closest to the most recent common ancestor of all scaly-foot snail populations, and it is thus treated as the outgroup for downstream gene flow analyses using SNPs (Figure 1c).

### Gene flow among groups

Significant gene flows (*Z* score >3 & *P* <0.01) were detected based on the *D*-statistics between: (1) Kairei-Solitaire and Wocan, (2) Kairei-Solitaire and Onnuri-Onnare, (3) Tiancheng and Onnuri-Onnare, and (4) Tiancheng and Longqi-Duanqiao (Figure 2a). In addition, the *f*-statistics revealed two pairs of gene flows with different intensities, with gene flow being more intense overall between two geographically closer populations. The gene flow between Kairei-Solitaire and Onnuri-Onnare was stronger than between Tiancheng and Onnuri-Onnare, and the gene flow between Onnuri-Onnare and Kairei-Solitaire was greater than that between Kairei-Solitaire and Wocan (Figure 2b).

**Figure 2.**
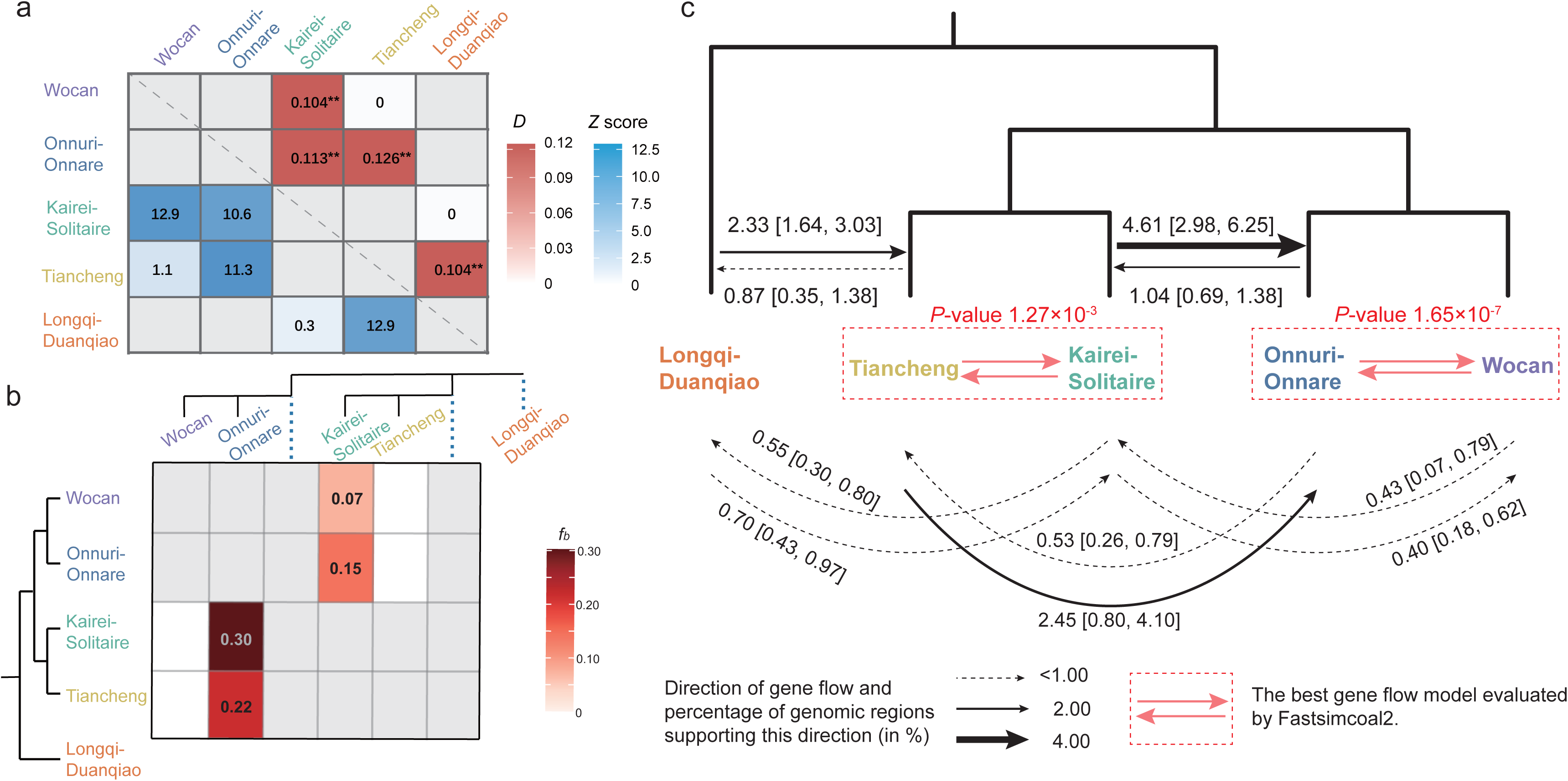
Phylogenetic relationships and gene flow between populations of the scaly-foot snail. **a.** Heatmap visualizing results of the ABBA-BABA (*D*-statistic) test for gene flow between populations. *D*-statistic values are represented by red blocks, and corresponding *Z*-scores (assessing significance of deviation from zero, indicating no gene flow) by blue blocks. A *Z* -score magnitude greater than 3 (*Z* score >3) indicates that the *D*-statistic is significantly different from zero. Asterisks (**) adjacent to *D* -statistic values denote statistical significance at *P* <0.01. Gray squares indicate population pairs that were not tested (sister species/populations). Values of 0 correspond to non-significant *D* -statistic results (*Z* score <3 or *P* ≥0.05). **b.** Heatmap of f-branch (*fb*) statistics for genetic populations using Dsuite. Calculation of *fb* statistics was constrained to the groupings of populations that fit the supplied population tree (see Figure 2b), which is shown along the y-axis. Each branch of the tree, including internal branches (e.g., blue dashed line representing an ancestral population), points to a corresponding row in the matrix with inferred *fb* statistics. The value in the matrix measures the extent of allele sharing between the corresponding branch of the population tree on the x-axis (relative to its sister branch) and the population on the y-axis (P3 in standard *D*-statistics). Gray cells indicate that calculation of *fb* statistics is not applicable given the population tree topology. For example, the highlighted (darkest) cell indicates significantly stronger allele sharing between the Onnuri-Onnare (x-axis) and the Kairei-Solitaire populations. c. *D*_FOIL_ test of gene flow direction among populations. The analysis was based on 2024 genomic regions generated from a 200 kb sliding window scan of the genome. Single-headed arrows indicate the inferred direction of gene flow. The numbers indicate the percentage (%) of genomic regions that support the corresponding gene flow direction. Dashed lines signify that gene flow between the two populations is supported by less than 1% of the genomic regions. To analyse the gene flow between sister groups using Fastsimcoal2, we pre-defined four gene flow models to fit the data. The optimal gene flow model was selected based on the Δlikelihood value, as indicated by the pink box and arrow in the figure, and the *P*-value was used to check the significance of the best-fit model versus the rest three models. The figure shows that the best-fit model for gene flow between the sister groups is continuous bidirectional (see Figure S8 for further details).

To determine the direction of gene flow, a *D*_FOIL_ test was used to detect distinct gene flow intensities on selected individuals in each genetic group (Figure 2c, Figure S7). The strongest signals occurred from Kairei-Solitaire to Onnuri-Onnare groups, which represented 4.61% of genomic regions on average. This was also highlighted by the presence of the hybrid-like individual (On8) in Onnuri, which exhibits the same direction of gene flow but represents an even higher proportion of genomic regions (8.3%) (Figure S7g). In contrast, the weakest was between Kairei-Solitaire and Wocan (0.40–0.43% of the genomic regions) (Figure 2c). Notably, the gene flow direction was consistently northwards across all populations. Three key examples illustrate this pattern: (1) from Longqi-Duanqiao to Tiancheng, contemporary gene flow was detected in 2.33% of the genomic regions (vs 0.87% in the reverse direction); (2) 4.61% from Kairei-Solitaire to Onnuri-Onnare vs 1.04% in the reverse direction; and (3) 2.45% from Tiancheng to Onnuri-Onnare, vs 0.53% in the reverse direction (Figure 2c).

To compensate for the limitations of *D*-statistics, *f*-statistics, and the *D*_FOIL_ test which can only detect and quantify gene flow outside sister populations, we used SNP simulations in Fastsimcoal2 - an approach that allows for a quantitative estimation of migration rates between adjacent populations, including pairs that could not be assessed by the *D*_FOIL_ test (Tiancheng and Kairei-Solitaire; Onnuri-Onnare and Wocan). We computed four distinct gene flow models (Figure S8) to identify the best-fit scenario. The continuous bidirectional gene flow model (including either symmetrical or asymmetrical) was selected as the best fit for all population pairs versus the other three models, i.e. No gene flow, Unidirectional (South to North), and Unidirectional (North to South) (*P* < 0.05)(Figure 2c), and the results were consistent with the *D*_FOIL_ test in the directionality of gene flow (Figure S7).

### Historical population demography

Based on phylogeny-based calculations (see Methods for details) with a high-quality genome of the closely-related confamilial snail *Gigantopelta aegis*, we estimated the genome-wide nucleotide substitution rate of the scaly-foot snail to be 7.9×10^-9^ [interquartile range: 6.1×10^-9^–1.1×10^-8^] per site per year following the neutral mutation theory, where the neutral nucleotide substitution rate is approximately equal to the mutation rate. We combined both PSMC (simulating 10,000-1,000,000 years ago) and popsizeABC (focusing on the last 10,000 years) methods to reconstruct the historical effective population size dynamics (Figure 3a & 3b). The PSMC results showed that the effective population size of Longqi-Duanqiao and other populations began to diverge between 200,000 and 400,000 years ago (Figure S9), which may correspond to when Longqi-Duanqiao split from the other populations. Between 100,000 and 200,000 years ago, the Wocan and Kairei-Solitaire populations began to diverge. We applied these divergence time ranges to the population history simulation of Fastsimcoal2 to estimate the average gene flow between populations. In addition, the population size fluctuation of Longqi-Duanqiao was smaller than other populations throughout history (Figure 3a; Figure S9).

**Figure 3.**
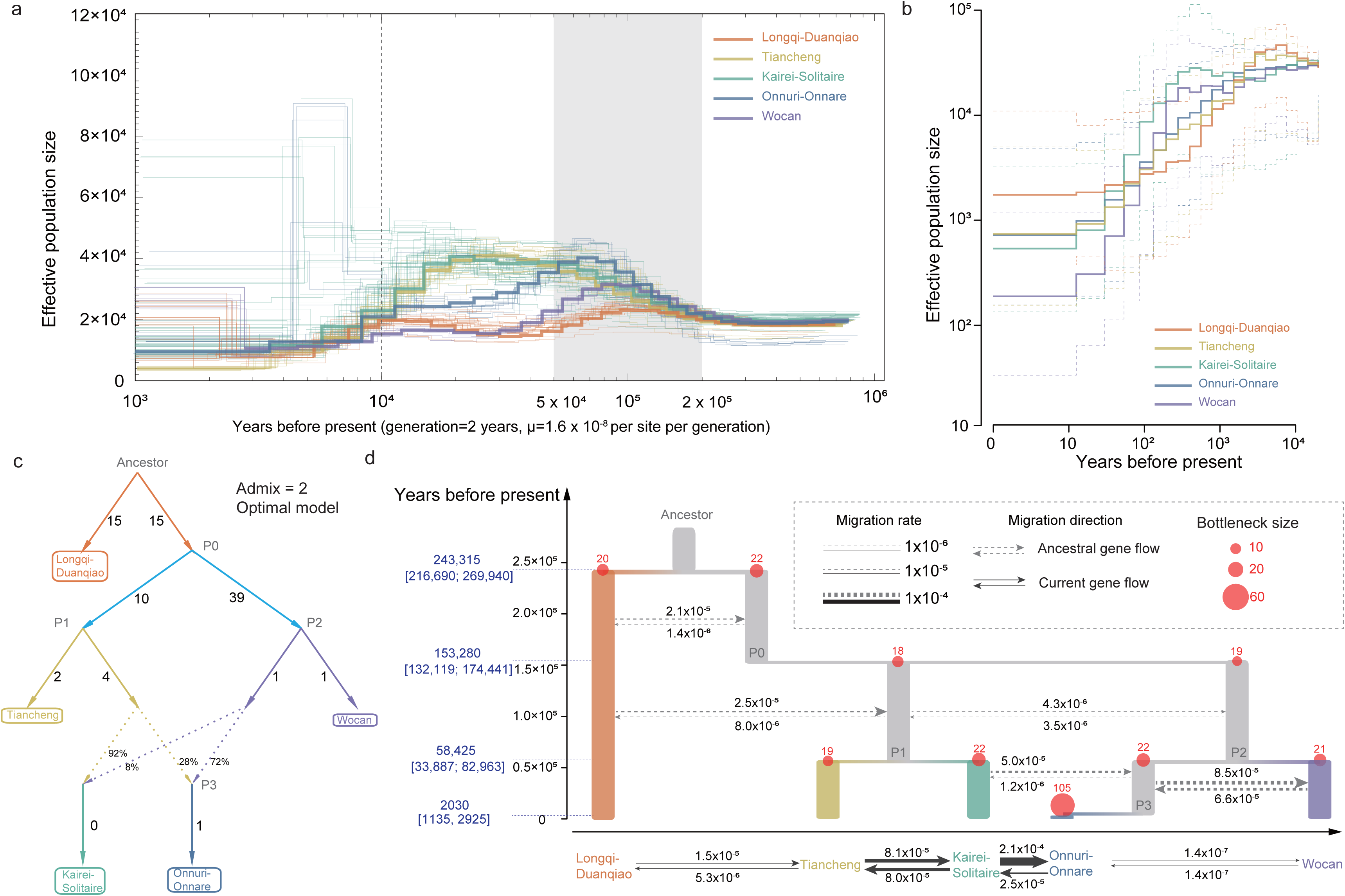
Demographic history inference of the scaly-foot snail. **a.** Long-term effective population size (*Ne*) trajectory of the scaly-foot snail estimated using PSMC (timeframe: ∼1 thousand years ago (Ka) to 1 million years ago (Ma)). PSMC has limited resolution for the recent past (<10 Ka). **b.** Recent *Ne* dynamics inferred using PopSizeABC (timeframe: present to ∼10 Ka). **c.** The highest-scoring demographic history and admixture graph with two admixture events inferred by qpGraph. Numbers on solid lines are inferred drift lengths. Percentages above dashed lines (admixture edges) show admixture proportions. The two ancient gene flow events include: (1) introgression from Kairei-Solitaire into the recent ancestor of Onnuri-Onnare, and (2) introgression from the recent ancestor of Onnuri-Onnare into Kairei-Solitaire. **d.** Population history model of the scaly-foot snail, modified from the optimal qpGraph model and constructed with population parameters estimated using Fastsimcoal2. In contrast to the pulse gene flow in the qpGraph model, this model incorporates continuous gene flow, considered more consistent with the species’ demographic history. The direction of gene flow is indicated by arrows: dashed arrows represent gene flow between ancestral populations (ancient gene flow), while solid arrows represent contemporary gene flow. Numbers on the arrows indicate the specific intensity of gene flow, expressed as the migration rate per year (m). Current populations are represented by differently coloured bars, while ghost populations are shown as grey bars. Red dots on the bars indicate population bottlenecks experienced at the time of population formation, with adjacent numbers representing the *Ne* during the bottleneck. The y-axis represents historical time in years before present, with numbers in parentheses indicating the 95% confidence intervals (CI). This analysis utilized a prior mutation rate range for the scaly-foot snail of [6.1×10^−9^ to 1.1×10^−8^] substitutions per site per year.

Our PopSizeABC results revealed that the effective population size of all populations has been declining during the past 10,000 years (Figure 3b). The largest effective population is Longqi-Duanqiao, while the smallest is Wocan; consistent with the field observations during sampling (Figure S10). scaly-foot snail is a dominant species in Longqi, but only a minor component in Wocan^18^. The linkage disequilibrium (LD) analysis showed that the LD decay rate of Longqi-Duanqiao was higher than Wocan, with the distance at which *r*^2^ decayed to 0.2 was 1025 bp for Wocan and only 38 bp for Longqi-Duanqiao (Figure S11). This further supports the current effective population size of Longqi-Duanqiao being higher than Wocan.

To explore the population demographic history, we also applied qpGraph to simulate the best population history model. Regardless of the number of mixing changes, the Longqi-Duanqiao population was the best fit as the outgroup data (Figure S5a, S5b, *P* <0.01). When admixture events (where previously isolated populations interbreed and form a new population with mixed ancestry) were 0, the formation of the population followed a stepping-stone model^38^ (Figure S5a). The formation process of the populations with the optimal model with the admixture events set to 2 was very complicated (Figure S5c), where, with the exception of Longqi-Duanqiao, the formation of all populations involved phantom populations (Figure 3c), also consistent with the results from badMixture (Figure S4).

We combined the optimal model obtained by qpGraph with the phylogenetic relationships among populations to construct a Fastsimocal2 historical model of the scaly-foot snail population demography that contained five phantom populations (Ancestor, P0-P3) (Figure 3d, Supplementary Data 4). The divergence time between Longqi-Duanqiao and the rest was around 243,315 [95% CI: 216,690–269,940] years ago, while the divergence time between Wocan and Kairei-Solitaire was around 153,280 [95% CI: 132,119–174,441] years ago, generally consistent with the aforementioned PSMC results. In addition, the divergence time between Kairei-Solitaire and Tiancheng to be around 58,425 [95% CI: 33,887-82,963] years ago, which roughly corresponded to previous estimates based on radioisotopes when hydrothermal activity began in Kairei (96,000 years) ^39^. Finally, the formation of Onnuri-Onnare and Wocan are both tightly linked with phatom populations P2 (and then P3 for Onnuri-Onnare), which explains why Onnuri-Onnare and Wocan exhibit a low degree of genetic differentiation (*P* <0.01) despite Kairei-Solitaire being geographically closer to Onnuri-Onnare.

### Physical ocean modelling

To understand the hydrodynamics transporting propagules in the study region, we tracked tracers released near the vent fields Longqi-Duanqiao, Tiancheng, Kairei, Solitaire, Onnuri-Onnare, and Wocan at a concentration of 1×10^6^ tracer units /km^2^ (released at a specific depth, confined to a horizontal plane) for 15 years (Longqi and Duanqiao, Onnuri and Onnare were grouped together due to limited geographic distances between them). Since the speed of deep ocean currents is very slow (a few centimetres per second) we chose a longer time interval of 15 years here in order to visualise the direction and extent of tracers exchanges among the vents, even though the scaly-foot snail larvae may not be able to survive for such a long time (however, the actual survival duration remains unknown). Limited by the accuracy of the ocean current model, the release depth of the numerical example (1500-3000 meters) cannot accurately correspond to the sampling depth (between 1780-2993 meters). The release depth is set as close as possible to the sampling depth of each vent field (the error is controlled within 1 km) in the bathypelagic zone (1000-4000 meters) (Figure 4).

**Figure 4.**
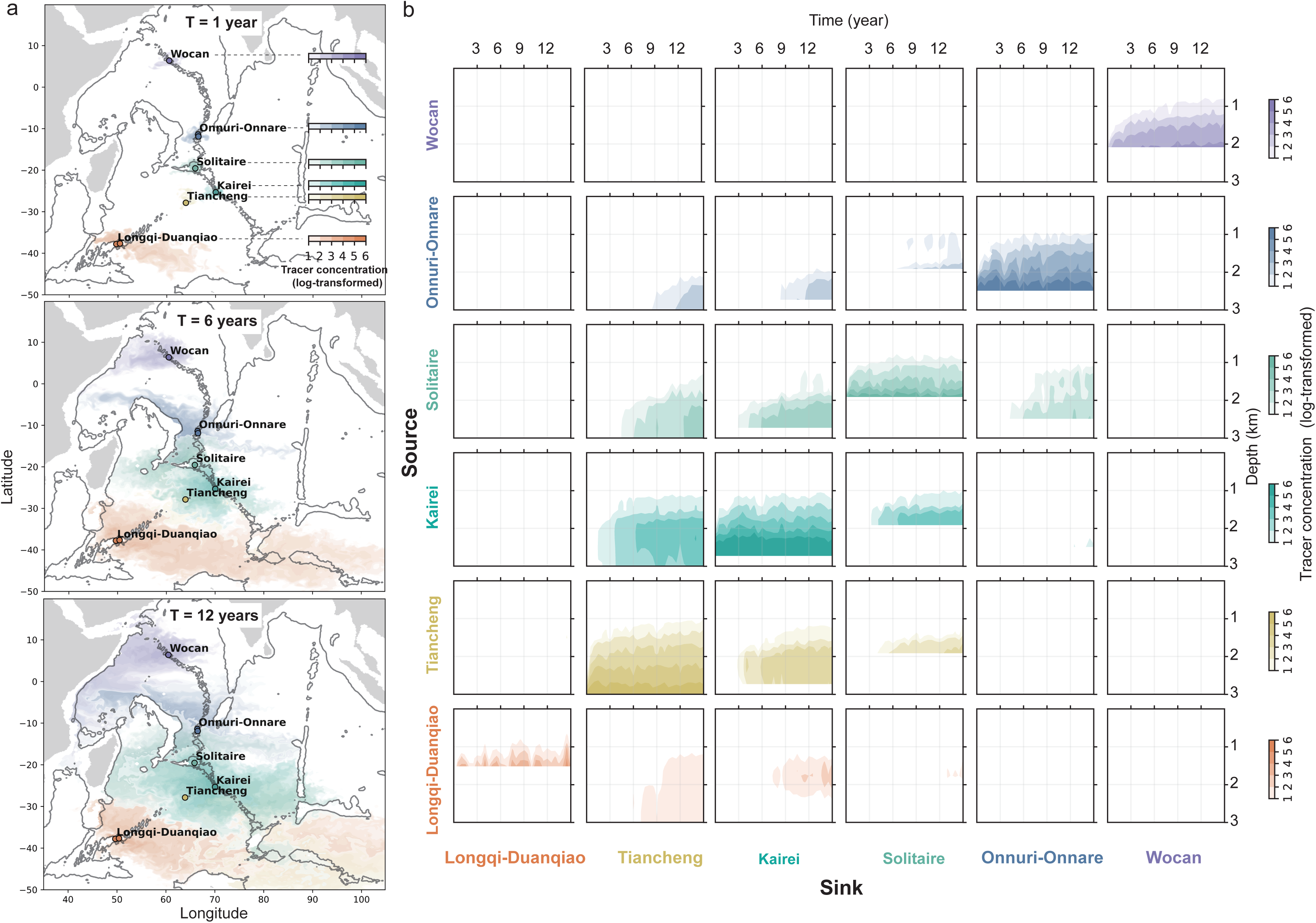
Tracers simulation of passive larval dispersal for the scaly-foot snail near Indian Ocean hydrothermal vents via ocean currents. **a.** Results from tracers simulation depicting the dispersal of scaly-foot snail larvae within deep Indian Ocean waters (1500– 3000 m depth) over the 1, 6, and 12 years. Tracers are color-coded by their source hydrothermal vent field: Longqi-Duanqiao (orange), Tiancheng (yellow), Kairei (cyan), Solitaire (green), Onnuri-Onnare (blue), and Wocan (purple). Color intensity represents the simulated larval particle density (log-transformed), as indicated by the color bar. **b.** Long-term larval connectivity matrix (15 years). Results from tracers drift simulation over 15 years, showing connectivity between different scaly-foot snail geographic populations (vent fields). The x-axis represents time (years), and the y-axis represents depth (km). Each stacked cell illustrates larval particles conveyed from a specific source population, labeled on the left of the panel, to a designated sink population, noted at the bottom of the panel. Tracer concentrations (log-transformed) is represented by colour intensity, using the same colour scheme and scale as in panel **(a)**.

In general, the tracers from Longqi-Duanqiao showed a wide dispersal range, the tracers of the three geographically intermediate vents (Tiancheng, Kairei, Solitaire) communicated closely, and the northernmost Wocan was isolated. The tracers released from Longqi-Duanqiao vent fields take up to seven years to reach Tiancheng with less than 0.001% of the tracers arriving, consistent with the gene flow order of magnitude (10^-5^) between Longqi-Duanqiao and Tiancheng from our fastasimcoal2 simulation. The average migration rate we obtained using the fastasimcoal2 simulation overlooks the complex population history between the two populations (such as phantom populations or population bottlenecks). The three intermediate release sites (Tiancheng, Kairei, and Solitaire) exhibited the most active tracer exchanges, with tracers reaching adjacent hydrothermal vents within three years at concentrations of up to 1% of the source release levels. Notably, Solitaire-originating tracers showed significantly greater delivery to Onnuri-Onnare (15-year period) than in the reverse direction, consistent with the asymmetric bidirectional gene flow between Solitaire and Onnuri-Onnare detected in genetic analyses.

## Discussion

### Genetic differentiation across vents

Our PCA and structure analyses identified five main genetic groups of scaly-foot snails: Longqi-Duanqiao, Tiancheng, Kairei-Solitaire, Onnuri-Onnare, and Wocan. These genomic findings offer much higher resolution than previous studies based solely on COI data^22, 35, 37^, and put forward these five groups as evolutionarily significant units (ESUs) with distinctive genetic diversity in a conservation context. The inclusion of the first genetic data of any kind from the Onnuri-Onnare scaly-foot snail population proved particularly revealing. Despite being relatively close to Solitaire, the Onnuri-Onnare group exhibited a higher genetic affinity with the distant Wocan population on the CR. This suggests complex historical or contemporary processes shaping its genetic makeup, agreeing with our demographic analyses revealing phantom populations being key in its formation. Furthermore, our findings reinforce the previous hypothesis^37^ that deep transformation faults on the ridges serve as key dispersal barriers for vent animals, explaining the high genetic differentiations seen between Longqi-Duanqiao and Tiancheng or between Kairei-Solitaire and Onnuri-Onnare. These probably also contribute to the formation of vent biogeographic provinces, separating sSWIR from nSWIR.

### Demography and phantom populations

Our data provides the first insights into the historical demography of Indian Ocean vent endemic species. Both mitogenome phylogeny and qpGraph simulations placed Longqi-Duanqiao as the earliest-diverging extant population, closest to the founding population of the scaly-foot snail in the Indian Ocean. The species appears to have subsequently spread northwards along the ridge all the way to Wocan. Our PSMC analysis indicates that Longqi-Duanqiao populations began to diverge from other populations between 200,000 and 400,000 years ago. Its historical fluctuation in effective population size was smaller than other populations, suggesting a relatively stable environment compared to other populations perhaps linked to the ultraslow spreading speed.

A key revelation from our demographic reconstructions, particularly from qpGraph and badMixture analyses, is the crucial role of phantom populations – either extinct historical populations or unsampled contemporary ones – in shaping the genetic landscape we observe today. The apparent genetic similarity between the geographically distant Onnuri-Onnare and Wocan populations, for instance, appears to be attributable to shared ancestry involving phantom populations (P2 and P3 in our model, Figure 3c and Figure S4) rather than significant contemporary gene flow. The three Indian Ocean ridges are known to host numerous inactive vent fields and unconfirmed hydrothermal plume signals^40^, which could represent these historical or even contemporary phantom stepping-stones. While pinpointing these phantom populations is challenging, it is clear that their importance is linked to the dynamic and often ephemeral nature of vent ecosystems over geological timescales^12^.

The estimated divergence times of around 58,425 [95% CI: 33,887–82,963] years ago between Kairei-Solitaire and Tiancheng, is similar to the estimated initiation of venting activity at Kairei hydrothermal vent activity initiation (96,000 years ago)^39^. While our demographic simulations posit that the Longqi-Duanqiao population has persisted for at least 216,690 years which is very long for vent fields, the longevity of vents on the mid-ocean ridges is linked to the spreading speed^12^ – SWIR is an ultra-slow spreading ridge which has been suggested to be capable of supporting stable venting activity at one field for such extended periods^41, 42^. Conversely, such long-term stability is very unlikely on intermediate-spreading ridges like the CIR or fast-spreading ridges like the well-studied East Pacific Rise^43, 44^. Furthermore, our PopSizeABC results indicate a decline in the effective population size across all populations over the past 10,000 years (Figure 3b), a trend that might be linked to sea-level fluctuations changes^45^.

### Gene flow patterns vs physical ocean modelling

The inferred contemporary and recent historical gene flow patterns, particularly the predominant south-to-north directionality (i.e., from Longqi-Duanqiao towards Tiancheng, and from Kairei-Solitaire towards Onnuri-Onnare and finally Wocan), align broadly with the general deep-water circulations as a part of the previously reported Great Ocean Conveyor Belt in the Indian Ocean^46, 47, 48^, which includes northward flows from the SWIR towards the Rodriguez Triple Junction and beyond. This is in line with the inferred larval dispersal mode of the scaly-foot snail, with lecithotrophic larvae carried by deep currents^35^.

Our physical ocean modelling experiments, simulating tracer dispersal over 15 years, lend support to these genomic reconstructions. The model showed limited tracer exchange between Longqi-Duanqiao and more northerly populations, consistent with the genetic isolation of Longqi-Duanqiao. The model also revealed closer connectivity among the central CIR sites (Kairei, Solitaire, Onnuri-Onnare), similar to results from the gene flow analyses. Notably, the physical ocean model predicted a higher probability of tracer transport from Solitaire to Onnuri-Onnare than in the opposite direction, matching the asymmetric gene flow detected. However, these simulations are subject to certain limitations. The 15-year tracer tracking period was necessary to observe any long-distance connections, but exceeds the likely PLD of lecithotrophic larvae estimated for peltospirid snails^15, 49^ which is thought to be up to several months. This suggests that successful long-distance dispersal in this context is likely a rare, multi-generational stepping-stone process, possibly involving unsampled contemporary phantom populations, rather than direct single-generation transport. We do note that since the larvae of the scaly-foot snail is entirely unknown, it may have longer PLDs than expected and requires future larval biology research to clarify. Furthermore, the resolution of our model – while at the state-of-the-art level for current global-scale simulations – may not capture all fine-scale bathymetric features or localized current dynamics critical for larval dispersal, especially given the general paucity of detailed oceanographic data for the deep Indian Ocean^50, 51, 52^. Notwithstanding these shortfalls, the congruence between genetic patterns, inferred gene flow, and modelled tracer dispersal together paints a compelling picture for how the scaly-foot snail (and likely other vent species with similar dispersal strategies) get around in the Indian Ocean.

### Priorities for conservation

All populations of the scaly-foot snail, except for Solitaire in the Exclusive Economic Zone (EEZ) of Mauritius, are located within ISA contract areas^18^ or is in the process of becoming one in the case of CR^24^. The ISA is currently in the process of drafting the Mining Code with the aim to finish the drafting process in 2025, which will set the standard and expected practice for how contractors proceed with future exploration and mining activities in the High Seas^53^. This overarching Code will include rules on environmental impact assessments, compliance and enforcement of regulations, benefit-sharing mechanisms, among other things – it will shape the landscape of deep-sea mining for the future decades to come. This timing makes our data especially crucial and timely for informing and guiding policymakers, the ISA, and other stakeholders including the general public about genetic structures and evolutionary significant units in Indian Ocean vents in order to inform conservation planning. As vent fields are typically small (much smaller than soccer fields), the removal of a vent site will almost certainly result in the extinction of the entire population there^20, 54^.

Our tracer release simulation showed that the direction of the SWIR deep current (from southwest to northeast) coincides with the direction of the unidirectional deep-sea thermohaline circulation. As such, Longqi-Duanqiao was not observed to receive larvae from other populations. The direction of dispersal from southern populations towards CR is consistent with the deep ocean current of the Great Ocean Conveyor Belt from south to north, and thus the Wocan population neither receives larvae from other populations nor contributes larvae, and as a result, is genetically isolated. It can be inferred that many other vent-endemic species living in Longqi-Duanqiao and Wocan vents that also employ deep-water dispersal using lecithotrophic larvae^35^, a common trait among vent species, may show similar dispersal patterns as the scaly-foot snail. These isolated populations harbour genetic diversity not found elsewhere, making them important evolutionarily significant units that may be in the early stages of speciation. As such, we propose Longqi-Duanqiao (within the Chinese contract area) and Wocan populations as the highest priority for protection. Similarly, Onnuri-Onnare (within the South Korean contract area) plays an essential role in maintaining the genetic diversity of the species and is also prioritised for protection.

Tiancheng, located in India’s contract area, is the only site with genetic connectivity to the Longqi-Duanqiao population through a weak gene flow, although simulations suggest that Longqi-Duanqiao could potentially also interact with Kairei within the German contract area. Perhaps it is possible to extract minerals from one of these intermediate sites without causing irreversible damage to the genetic diversity of scaly-foot snails, but a significant complication is that all these sites are under different contractors. In this case, the conservation effort and management plan have to be internationally orchestrated and decision-making. Such coordination is not required under the current contractual framework, and the ISA should play a central role in facilitating such efforts among contractors for a responsible and sustainable path forward.

## Conclusion

Our population genomics study of the iconic and ironclad scaly-foot snail included 125 individuals from eight distinct vents, providing an unprecedented level of insights on the connectivity and demography for an Indian Ocean vent-endemic animal. Our results highlight the key role of previously unrecognised phantom populations in shaping the genetic structure and diversity we see today. Our data shows a predominantly northward gene flow across all populations, and also supports deep transformation faults as important dispersal barriers limiting gene flow in the scaly-foot snail and likely other deep-dispersing species. Together, these provide novel insights into how the three currently recognized biogeographic provinces of Indian Ocean hydrothermal vents may have formed, and further reveal the potential for Onnuri-Onnare to house genetically distinct populations from other CIR sites. We propose the earliest-diverging source population Longqi-Duanqiao and the highly isolated Wocan as two critical sites to prioritise for protection in future conservation and management plans; though the five main genetic groups identified may all be considered evolutionarily significant units.

Many Indian Ocean vent species have been assessed as Endangered or even Critically Endangered on the IUCN Red List due to upcoming mining ^20, 55^, and our new data on the scaly-foot snail also helps inform the conservation of many other such vent-endemics with similar dispersal modes. Even with the additional recently discovered populations of the scaly-foot snail sampled herein increasing its known distribution, a reassessment would still result in it being considered threatened with extinction under the Vulnerable category, given there are less than 10 locations and all except Solitaire are under mining licenses^54^. Furthermore, the population genomics data presented here sets a standard for management of vents based on ESUs of genetic diversity within a species, which is not sufficiently reflected in the species-based IUCN Red List^56^ but also a crucial aspect of biodiversity. Given the iconic standing and a broad distribution, the scaly-foot snail can be a great example of flagship and umbrella species in deep-sea conservation – protecting this fantastic beastie will also lead to the survival of many others. Coordinating conservation efforts presents a special difficulty in the Indian Ocean as almost every population is under a different mining contractor with the ISA. Therefore, we recommend the ISA to oversee and mediate a framework for concerted international collaboration among the contractors or a moratorium until this can be achieved, when finalising the Mining Code.

## Materials and Methods

### Sample collection and genomic DNA extraction

A total of 125 scaly-foot snails (*Chrysomallon squamiferum*) were collected from eight hydrothermal vent fields on the Indian Ocean Ridges from 12 research cruises between 2013 and 2023 (Figure 1a; Table 1). Sampling sites spanned three mid-ocean ridges: SWIR (Longqi, Duanqiao, and Tiancheng), CIR (Kairei, Solitaire, Onnuri, and Onnare), and CR (Wocan). After recovery on-board the ship, snails were either immediately preserved in liquid nitrogen and stored at -80°C, or preserved in 95% ethanol for further usage. All transportations of frozen material were done on dry ice.

For DNA extraction, the foot tissues (∼500 mg) were thawed and extracted using DNeasy Blood & Tissue Kits (Qiagen, Germany). DNA quality and integrity were assessed via NanoDrop spectrophotometry, agarose gel electrophoresis, and Qubit fluorometry. Sequencing libraries (350 bp insert size) were prepared from purified DNA using the NEBNext Ultra II DNA Library Prep Kit (New England Biolabs, USA) for sequencing on the Illumina HiSeq X Ten (Illumina, USA), NovaSeq 6000 (Illumina, USA) or NovaSeq X Plus platform (Illumina, USA) (paired-end mode and 150 bp read length) (for details, see Supplementary Data 1).

### Genome-wide SNP calling

Raw reads were filtered for adapters and low-quality sequences using fastp v0.23.4 ^57^ with the settings of *--length_required 60 --qualified_quality_phred 20 --unqualified_percent_limit 20 --n_base_limit 5*. High-quality reads were mapped to the *C. squamiferum* reference genome ^58^ using bowtie2 v2.5.4 ^59^ with the settings of *--sensitive-local*. Resultant SAM files were converted to sorted BAM files using SAMtools v1.20 ^60^. Duplicate mapping from polymerase chain reaction was marked using *MarkDuplicates* module of GATK v.4.5.0.0 ^61^. Sample-specific read groups were assigned using the *AddOrReplaceReadGroups* module in GATK.

To obtain high-quality SNPs, SNP calling was performed using two pipelines (i.e. GATK and BCFtools). In the GATK pipeline, SNPs were initially called for each individual using *HaplotypeCaller* in GVCF mode. The resulting GVCF files were then merged via *CombineGVCFs*, followed by joint genotyping with *GenotypeGVCFs* to generate population-level SNPs. For SNPs filtering, the whole procedure followed the GATK Best Practices Workflows (https://gatk.broadinstitute.org/hc/en-us/articles/360035890471-Hard-filtering-germline-short-variants). (1) The SNPs were filtered with the following settings: *QualByDepth (QD) < 2.0*, *FisherStrand (FS) > 60.0, RMSMappingQuality (MQ) < 40.0, MQRankSum < −12.5*, *ReadPosRankSum < −8.0,* and *StrandOddsRatio (SOR) > 3.0.*( 2 ) For each SNPs, we applied a sequencing depth (DP) filtering, and only positions with a *DP > 0.5× mean depth* and *DP < 2× mean depth* (*mean depth* represents the average depth of all samples, which is 29.52*×* in this study, see Supplementary Data 1). This strict DP filtering minimized the effects of sequencing errors in regions of low sequencing coverage and multiple-mapping errors in high-coverage regions ^62^. (3) For genotype quality (GQ) filtering, to ensure high-confidence genotype calls, we removed sites had a *GQ < 60* (GQ ≥ 60 corresponds to a 99.9999% confidence level in genotype accuracy).

In the BCFtools pipeline, BCFtools v.1.20 was used to call SNPs based on the combination of all the *.bam* files of all *C. squamiferum* samples. The “*--annotate*” flag in “BCFtools mpileup” and the “*--multiallelic-caller*” and “*--group-samples*” flag in “BCFtools call” were applied to call SNPs. “*--group-samples*” flag can reduce the number of false-positive callings in low-coverage samples. The filtering settings for DP and GQ steps are the same as the above GATK pipeline. Multiallelic sites were excluded in genotype calling.

To increase confidence in rare allele identification and eliminate potential somatic mutations, allelic balance filtering was performed for SNPs containing a single minor allele in 125 individuals. This filtering process was executed using allelicBalance.py (https://github.com/joanam/scripts/blob/master/allelicBalance.py). Only SNPs that were identified by both pipelines were retained for the downstream analyses.

Two distinct SNP Datasets were prepared for downstream analyses according to specific analytical requirements. For Dataset 1, filtration through VCFtools v.0.1.17 ^63^ was conducted with the following criteria: (1) only biallelic SNPs were retained (--*min-alleles* 2 --*max-alleles* 2); (2) sites present in ≥80% of 125 individuals (--*max-missing* 0.8); and (3) global minor allele frequency (MAF) ≥0.05 (*--maf 0.05*). For the purpose of investigating population history dynamics, Dataset 2 was processed without MAF filtering (*--maf 0*), thereby retaining all rare alleles in the analysis ^64^.

### Population structure analysis

For Dataset 1, linkage disequilibrium (LD) pruning was used to remove SNPs that are in high LD to ensure that the SNPs being analysed are independent. Data pruning was achieved using the *--indep-pairwise 10kb 1 0.2* parameter in PLINK v.2.00 alpha ^65^. Subsequently, principal component analysis (PCA) was conducted on all 125 samples of scaly-foot snails with the *-- pca* parameter in PLINK. The population structure and individual ancestry were then examined using the maximum likelihood (ML) estimation method implemented in ADMIXTURE v.1.3.0 ^66^. The number of genetic groups, represented by *K*, was predefined to range from 2 to 8. To evaluate the optimal number of genetic groups that best fit the data, the cross-validation (CV) procedure in ADMIXTURE was applied with the *--cv* flag.

To quantify the extent of genetic differentiation among populations of scaly-foot snails, Hudson’s genome-wide *F*_ST_ estimates were calculated based on Dataset 1. This analysis was performed using PLINK v2.00 alpha with the parameters *--fst CATPHENO method=Hudson blocksize=1000000*.

### Determining the first diverging population closer to the ancestral population

To identify which of the sampled populations is genetically closest to the ancestral population, we employed three distinct methodologies: (1) constructed a phylogenetic tree using mitochondrial coding genes from scaly-foot snails with the closely related neomphaline snail *Cyathermia naticoides* (NCBI accession No.: BK064858.1) as the outgroup. We assembled 125 scaly-foot snail mitochondrial genomes using NOVOPlasty v4.3.5 ^67^ with the following settings: *Genome Range = 18000-24000, seed =* Chrysomallon squamiferum *Coding Sequences file (NCBI accession number: AP013032), K-mer = 33, Read Length = 150, Insert size auto = yes, and Use Quality Scores = no*. Subsequently, we extracted all coding genes from all scaly-foot snail samples using BLASTn v2.16.0 with the parameter *- outfmt 6* and a FASTA file of scaly-foot snail mitochondrial coding genes ^68^. Multiple sequence alignment was performed using MAFFT v7.526 with the *--auto* option. Finally, a partitioned phylogenetic tree was constructed using IQ-TREE 2 v2.3.6 ^69^ with the parameters *-p partition.nex -B 1000 -m MFP*. (2) To investigate recent demographic history, we implemented badMIXTURE, an approach that assesses the goodness of fit of admixture models using ancestry “palettes” estimated by CHROMOPAINTER. Our analysis followed the example script provided by the software authors (https://github.com/danjlawson/badMIXTUREexample). (3) Outgroup Selection and Model Fitting: We utilized qpGraph ^70^ and Fastsimcoal2 ^71^ software to determine the optimal outgroup. This involved constructing various models by systematically rotating different genetic populations as the outgroup. The model exhibiting the highest goodness-of-fit score, indicating the best fit to the data, was selected as the optimal model. This, in turn, identified the most suitable outgroup. Details of the constructed models are provided in Figures S5 & S6, and the model files are publicly available on GitHub. Fastsimcoal2 was run with the following settings: *-m -0 -C 10 -n 100000 -L 40 -s 0 -M -c 0 -q*.

### Phylogenetic tree construction

For the second method, a p-distance matrix was calculated using VCF2Dis v.1.52m ^72^ based on the Variant Call Format (VCF) input. The data were repeatedly sampled 1000 times with the *-Rand 0.25 -TreeMethod 1* settings, and the resulting trees were constructed using the “*fneighbor*” module in the PHYLIPNEW-3.69.650 software. Finally, all trees were merged, and bootstrap values were calculated using the “*percentageboostrapTree.pl*” script in the VCF2Dis tool.

### Gene flow investigation and demographic reconstruction

To verify whether there is gene flow between different scaly-foot snail populations and the relative strength of gene flow between populations, Dsuite v.0.5 r52^73^ and Dataset 2 with rare alleles retained were employed to calculate Patterson’s D with *Dtrios* module and f4-ratio with *Fbranch* module. Patterson’s D, also known as ABBA-BABA, and the related estimate of admixture fraction *f*, referred to as the f4-ratio are commonly used to assess evidence of gene flow between populations in genomic datasets^73^. Based on the conclusions in the chapter “Determining the first diverging population closer to the ancestral population”, the Longqi-Duanqiao group was used as the outgroup to verify the gene flow between other populations. Calculations were limited to the groupings of populations that fit with a supplied tree based on *p*-distances calculated by IQ-TREE and VCF2Dis.

To verify the direction of gene flow, The *D*_FOIL_ tool^74^ was employed to perform Introgression Testing for the identified genetic groups. This tool only requires one individual for each population. Based on the phylogenetic tree constructed with VCF2Dis, samples that fit the required *D*_FOIL_ phylogenetic relationship of (((P1, P2), (P3, P4)), Po) were selected for analysis. Gene flow on the same branch (i.e., between P1 and P2, and between P3 and P4) cannot be tested by *D*_FOIL_ ^74^, therefore, Fastsimcoal2 analysis was additionally introduced. A total of four different gene flow models were constructed. The models are categorized based on the directionality of gene flow: “ongoing gene flow” (continuous bidirectional, symmetric or asymmetric, gene flow since population divergence), “no gene flow” (no gene flow occurs following the population split), and “Sou2Nor” and “Nor2Sou” gene flow (unidirectional gene flow since population divergence). Fastsimcoal2 was utilized to identify the optimal gene flow model among the identified genetic groups of scaly-foot snails.

The specific analysis method is as follows: the sorted and filtered BAM files corresponding to two individuals with the highest sequencing depth for each population were selected. To generate the consensus sequence for each population, ANGSD v.0.940-stable ^75^ was used with the parameters *-doFasta 2 -doCounts 1*. The genome was subsequently divided into 200 kb windows using seqkit. Windows located in the same genomic regions for each penta-taxon were then merged and filtered, discarding those with an *N* proportion greater than 50%.

Genotype statistics for the divided windows were computed using the *fasta2dfoil.py* script from the *D*_FOIL_ tool, followed by the application of the *dfoil.py* script to calculate gene flow between populations.

### Deducing the mutation rate of the scaly-foot snail

The nucleotide mutation rate estimation was conducted by referring to the method described by Cui et al.^76, 77^. The specific calculation method is as follows: To obtain the ratio of nucleotide substitution rates at neutral sites between scaly-foot snails and closely related species. Comparative genomic analyses were performed using reference genomes from eight gastropod species (Supplementary Data 5). Whole-genome coding sequences (CDS) and protein sequences (n = 197,701) were processed with OrthoFinder v2.5.5 ^78^ to identify 3112 universal single-copy orthogroups. These protein sequences were aligned using MAFFT v7.520 ^79^ with default parameters and subsequently converted to codon-based nucleotide alignments using PAL2NAL v14.1 ^80^. Concatenated codon-aligned sequences (1,822,122 bp) were partitioned by codon positions for model selection in IQ-TREE v2.2.2.7 ^69^ under “-m MFP+MERGE -B 1000” parameters. Optimal substitution models were identified as GTR+F0+R3 (codon position 1) and GTR+F0+R6 (positions 2-3). RAxML-NG v1.2.1 ^81^ was employed to refine branch lengths using 352,908 quadruple degenerate sites, yielding branch lengths of 0.8407 (*C. squamiferum*) and 1.0862 (*Gigantopelta aegis*).

The mutation rate is calculated based on the number of substitutions (*K*s) of synonymous substitution sites in all single-copy homologous genes and the divergence time between species. From 9162 orthologous gene pairs, ParaAT v2.0 ^82^ generated codon alignments for *K*a/*K*s analysis using KaKs_Calculator v3.0 ^83^ with Muse-Gaut model (AIC-selected). To obtain more reliable conclusions in subsequent population history simulations, we strictly filtered homologous genes. Only homologous gene pairs with values *P* < 0.01 (Fisher’s Exact test), and length greater than 300 bp were retained. Based on the calculation and statistics of 6025 homologous gene pairs, we used the upper quartile (Q3) per site/per year and the lower quartile (Q1) per site/per year as the range of mutation rates. mutation rates were calculated as:

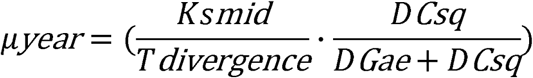

Where *D_Csq_* represents branch lengths of *C. squamiferum*. *D_Gae_* represents branch lengths of *G. aegis*. *T_divergence_* represents divergence time between *C. squamiferum* and *G. aegis*, which is 116 million years ago ^84^. *Ks_mid_* represents the median *Ks* value of 6025 homologous genes.

### Demographic history reconstruction

The PSMC method ^85^ was employed to estimate the effective population size (*Ne*). The analysis followed the best practices suggested by the software developers, with slight adjustments due to software version changes: The *-C50* flag in the BCFtools *mpileup* module and the *-c* flag in the BCFtools *call* module were used to call genotypes for the BAM file of each individual. All variable and non-variable sites were retained, and then *vcfutils.pl* program in the PSMC package was used to mask sites with depths less than one-third or greater than twice the average depth. The filtered VCF output was converted to FASTQ format. The “*fa2psmcfa*” and “*splitfa*” programs in the PSMC package were used to generate the required input for PSMC and split long sequences into short fragments. Finally, the rest of the parameters of the psmc module (*-N30 -t15 -r5 -p “4+25*2+4+6"*) were kept the same as the developer’s suggestions. The analysis was plotted using the Perl script *psmc_plot.pl* in the psmc package. A mutation rate of 6.1 × 10⁻^9^ to 1.1 × 10⁻⁸ per site per year (estimated in the previous section) was applied, along with an assumed generation time of two years for scaly-foot snails ^86^. For the 125 samples, we excluded samples with a sequencing depth less than 10**×** (sample number: WC2) or greater than 60**×** (sample numbers: WCB01, KRSFGB1, Na10). The remaining 121 samples were summarized in one figure and visualized using Adobe Illustrator CC. Different groups were distinguished by different colours, and the curve of one of the samples was bolded (Figure 3a).

As the PSMC method has a low resolution for recent history, PopSizeABC^87^ was used to infer recent changes in population size. Average LD and folded site spectrum data were constructed for each population across different physical distances, using a Dataset 1 that retained only common variants. PopSizeABC was executed to estimate effective population sizes over time within an Approximate Bayesian Computation (ABC) framework ^87^. The tool was run with the software developers’ recommended parameters, performing 200,000 simulations across 100 bins of 2 Mb each. A minimum MAF threshold of 0.2 was applied to both the site spectrum estimates and the LD calculations. It should be noted that we modified the author’s simul_data.py script so that the mutation rate is no longer a fixed value. We used the prior distribution range of scaly-foot snails. The modified script has been uploaded to GitHub.

To verify the demographic history of the scaly-foot snail, qpGraph analysis was performed using the *R* package ADMIXTOOLS2^70^. Following this, the population history was simulated using the model-based composite likelihood method implemented in Fastsimcoal v.2.8 ^71^, based on Dataset 2 with all rare variant sites retained. The two-dimensional Minor Allele Frequency Site/Allele Frequency Spectrum (MAF_SFS) was constructed using easySFS v.0.0.1^88^. The first step was to determine the projection value of each population using the *-- preview* setting. The second step was to specify the projection value of each population according to the number of maximizing separation sites with the settings *-a --proj 108,48,32,40,18 --order kairei_solitaire,longqi,onnuri_onnare,tiancheng,wocan --dtype int*. A total of 10 SFS files were generated between each of the five lineages.

Then, differentiation patterns, *Ne*, divergence time, and gene flow were simulated by performing 100 runs, 40 expectation maximization cycles, and 100,000 coalescent simulations for each model in Fastsimcoal v.2.8 with the settings “*-m -0 -C 10 -n 100000 -L 40 -s 0 -M -c 0 -q”*. Again, we used the range of the prior distribution (5 × 10⁻^9^ to 2 × 10⁻⁸ per site/per year) of mutation rates for this analysis. In addition, we used a priori inbreeding coefficients for each population (Figure S12, Supplementary Data 2). Model goodness of fit was assessed by simulating SFSs for each demographic model and comparing the simulated likelihoods with the observed likelihoods. All model files (*.tpl* and *.est* file) are available in GitHub. The best demographic model, *N*_e_, and gene flow were determined based on the minimum delta likelihood, Akaike’s information criterion (AIC), and AIC weights. To obtain 95% confidence intervals for the best model, 50 new SFSs were generated by bootstrapping using ANGSD and parameter simulations were performed with the same number of runs, replicates, and simulations.

### Physical ocean modelling

We used the Massachusetts Institute of Technology general circulation model (MITgcm)^89,90,91^ to conduct a global ocean-sea ice coupled simulation. The model has a horizontal grid resolution of 1/6° and 63 vertical layers. The horizontal grid used is called LLC540 (Lat-Lon-Cap 540), which is a grid-refined version of the Estimating the Circulation and Climate of the Ocean (ECCO) version 4 grid and thus permits to resolve the relatively large ocean mesoscale activities, which can play important roles in the material transports even in the deep ocean. The model grid consists of five faces implemented with 13 tiles^92^. The vertical grid spacing ranges from 10 metres near the surface to 457 metres near the ocean bottom.

Forcing data for the model is obtained from the climatological annual cycle (averaged from 2000 to 2020) of the fifth generation ECMWF atmospheric reanalysis products (ERA5). This includes variables such as the 2 metre air temperature, specific humidity, precipitation, downward long/short-wave radiation, 10 metre zonal/meridional wind speed, and runoff. The model employs minimal parameterization, with the K-profile parametrization used for the surface boundary layer. A quadratic bottom drag with a coefficient of 0.0021 is applied. Additionally, the vertical diffusivity of temperature and salinity varies with latitude in this model.

To investigate the transport dynamics driven by large-scale ocean gyre circulations and mesoscale activities, six passive tracers were deployed at the ocean bottom of six hydrothermal vent stations: Longqi-Duanqiao, Tiancheng, Kairei, Solitaire, Onnuri-Onnare and Wocan. After a 50-year model spin-up, the tracers were introduced with no explicit diffusivity. Consequently the tracer transport is governed solely by resolved ocean currents and mesoscale activities. This setup mimics the passive drift of entities, such as certain deep-sea organisms, that are advected by ambient flows without active movement.By using passive tracers in this way, we effectively simulate the behavior of an extremely large number of Lagrangian particles transported by ocean circulation.

The tracer dynamics are described by the equation:

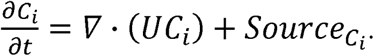

Where C_i_ represents the tracer concentration at each station, U denotes the three-dimensional velocity field, and Source is the initial tracer concentration at the simulation’s onset. To maintain sufficient tracer presence for robust transport analyses, concentrations at each station are relaxed to their initial values with a time scale of 86,400 seconds (i.e., one day). This strong relaxation, while not reflective of continuous biological presence, optimises computational efficiency by ensuring sustained tracer availability for transport across diverse ocean regions.

The tracers evolved over 15 years, with snapshots of their distributions recorded every six months. This configuration enables a detailed examination of long-term tracer dispersion driven by the ocean circulation, providing insights into connectivity and transport pathways in the deep ocean.

## Data availability

The Illumina raw sequencing reads were deposited in the NCBI SRA database under accession code PRJNA1270171.

## Code availability

All commands, codes, and intermediate files for the bioinformatics analyses are based on the available software listed in the ‘Methods’ section, which are provided at https://github.com/ChenXXXi/Population-genetics-of-Scaly-foot-Snails under the GNU General Public License v3.0.

## Acknowledgement

This study was financially supported by National Natural Science Foundation of China (No. 42176110), the Science and Technology Innovation Project of Laoshan Laboratory (LSKJ202203104), Natural Science Foundation of Shandong Province (ZR2023JQ014), the grant from Natural Science of China (HJRC2023001), the PI project (2021HJ01) from the Southern Marine Science and Engineering Guangdong Laboratory (Guangzhou). This research was also supported by a project titled ‘Understanding the deep-sea biosphere on seafloor hydrothermal vents in the Indian Ridge (No. 20170411)’ funded by the Ministry of Oceans and Fisheries, Korea. We thank the captain, chief scientist and crew member during the cruises of DY34, DY35, DY38, DY52-III, YK13-02, YK13-03, YK16-02E, and KIOST, and extend the same to the submersible technical team and pilots including those of HOV *Jiaolong*, HOV *Shinkai 6500*, ROV *ROPOS*, and ROV *Jason I*. We thank Dass Bissessur (Maritime Zones Administration & Exploration, Ministry of Defence and Rodrigues, Mauritius) for his help in obtaining permission to sample in the Mauritian EEZ, which was approved by the Ministry of Foreign Affairs, Regional Integration, and International Trade, Mauritian Government under ref. 29/2014; 50/38/24 V2. We thank Chenggang Liu for providing the photograph of the scaly-foot snail collected from Tiancheng. The bioinformatic analyses were performed on the High-Performance Biological Supercomputing Centre at the Ocean University of China. The physical ocean modelling simulations were carried out using the high-performance computing resources of the High-end Computing Centre at the Jiangwan Campus of Fudan University.

