## Supplementary Figures for "Population genomics of the endangered scaly-foot snail defines conservation units amid deep-sea mining threats in Indian Ocean vents"

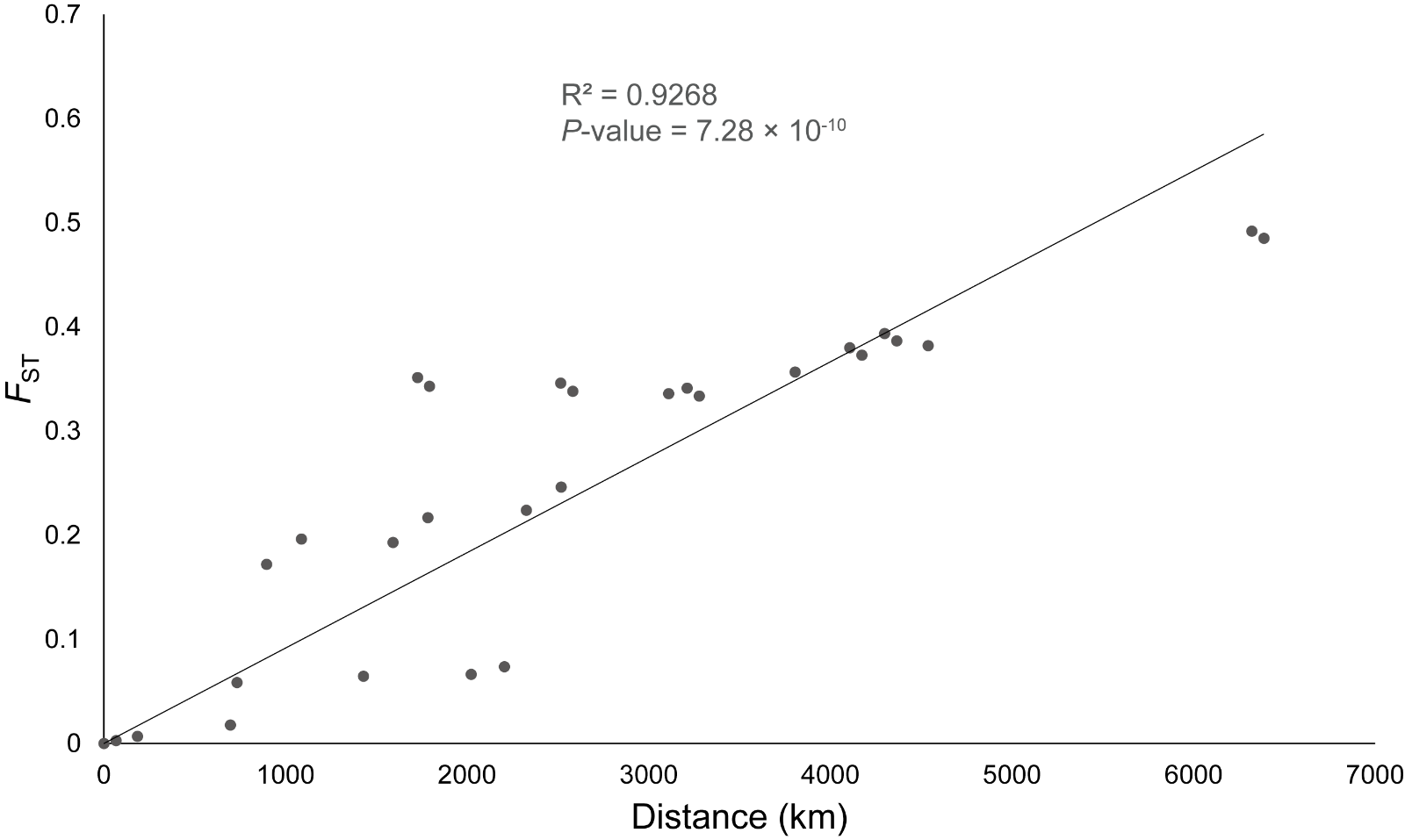

**Figure S1. Linear regression plot of *F*_ST_ values between two populations of the scaly-foot snail and geographical distance along mid-ocean ridges of the Indian Ocean.** The x-axis is the geographical distance and the y-axis is the *F*_ST_ value. The *P*-value is marked in the figure and is much less than 0.01, indicating that geographical distance is significantly correlated with *F*_ST_.

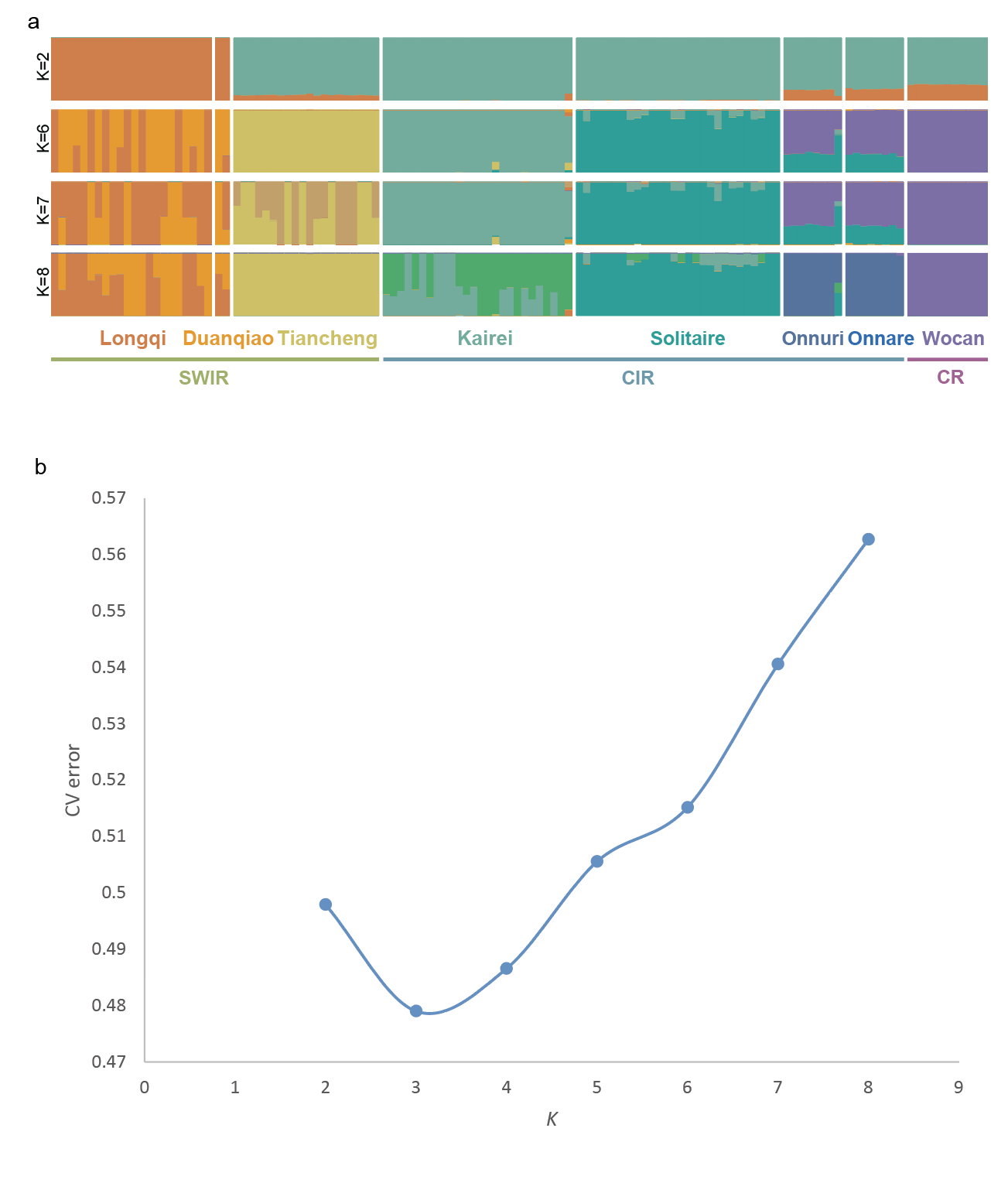

**Figure 2. Population structure of the scaly-foot snail revealed by ADMIXTURE analysis. a** The inferred ancestry for individuals assuming *K*=2, 6, 7, and 8 ancestral populations. Each vertical bar is an individual, colored by its proportion of ancestry from each cluster. Southwest Indian (SWIR), Central Indian (CIR), and Carlsberg (CR) ridges. **b** The accompanying plot shows the cross-validation error for each value of *K*, with the lowest error value at K=3.

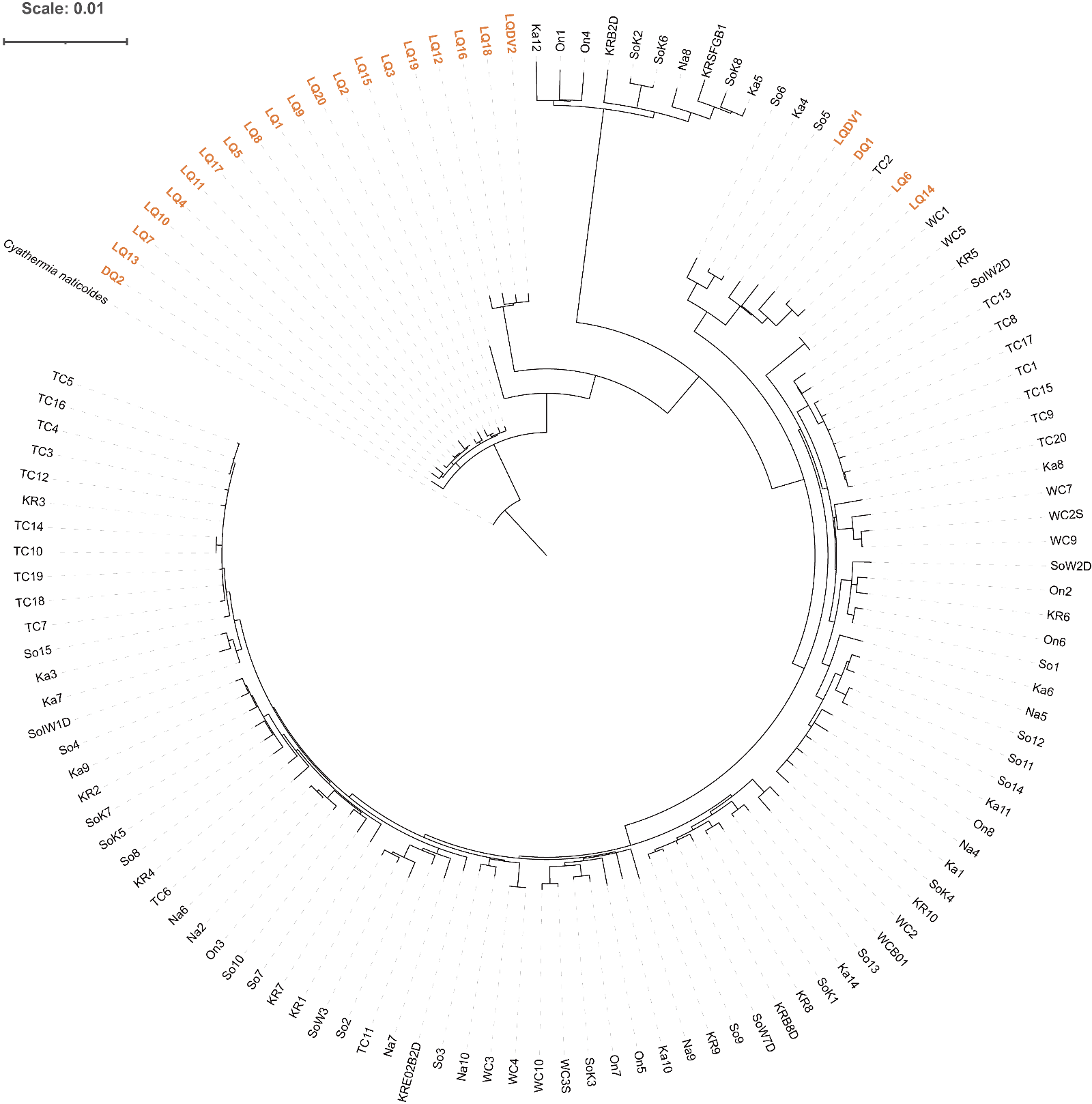

**Figure S3. Phylogenetic tree of the scaly-foot snail based on the concatenated sequences of 125 mitochondrial protein-coding genes.** The tree was constructed using the Maximum Likelihood method and rooted with the neomphaline snail *Cyathermia naticoides* as the outgroup. The Longqi-Duanqiao group show the closest phylogenetic affinity to the *C. naticoides*, suggesting Longqi-Duanqiao represents the earliest-diverging group compared to the others.

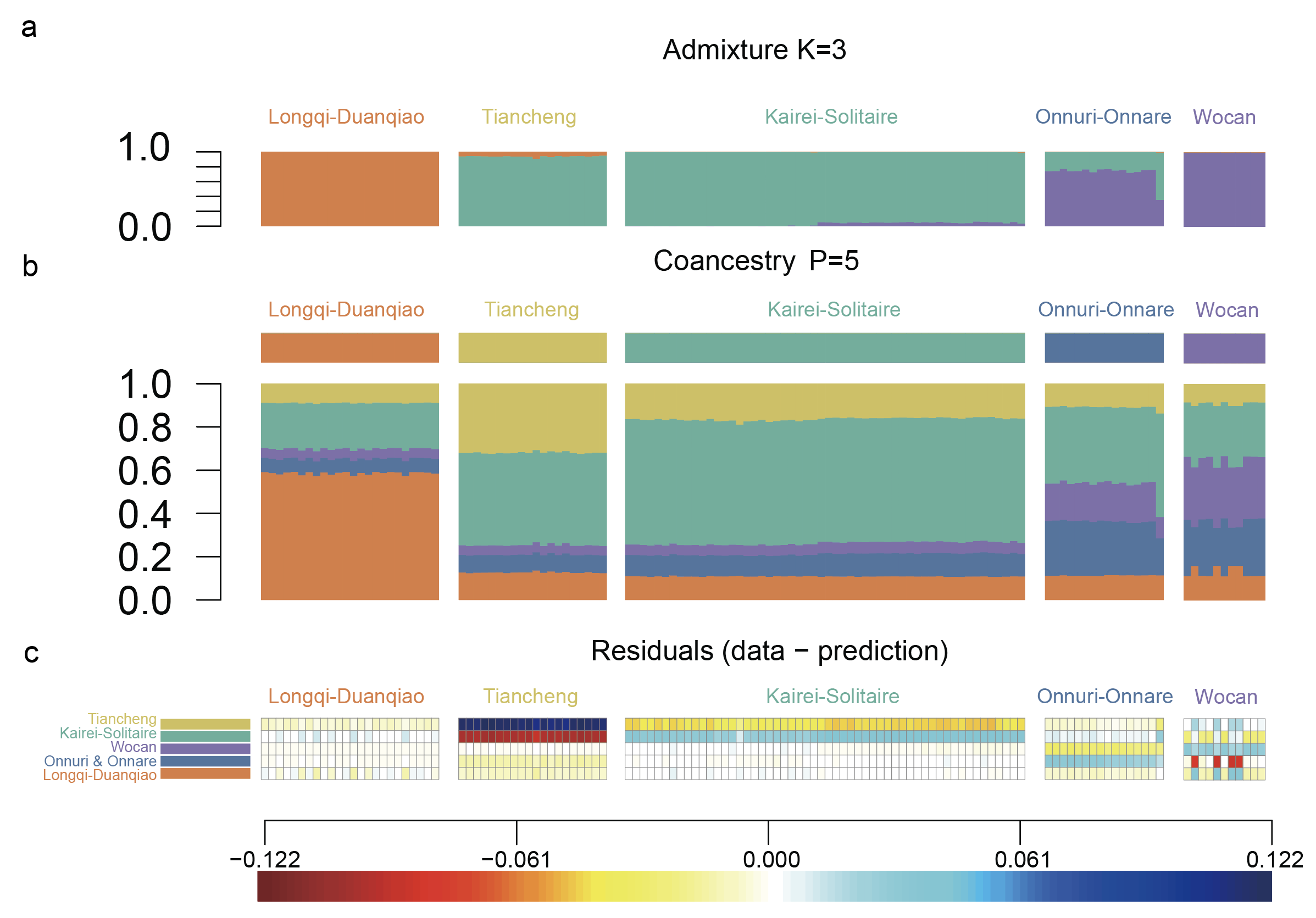

**Figure S4. The results of the badMIXTURE analysis showed that the scaly-foot snail of Longqi-Duanqiao contained the ancestry of all other groups. a** The inferred ADMIXTURE plots at K = 3. **b** The CHROMOPAINTER inferred painting palettes. **c** The painting residuals after fitting optimal ancestral palettes using badMIXTURE, presented on the residual scale shown. The presence of structured, non-random residuals for all populations, excluding Longqi-Duanqiao, reveals that the Admixture K=3 model (panel a) is a poor fit to the data (a "bad mixture"). This implies that the ancestry proportions shown in panel (a) should not be interpreted literally, as doing so could be misleading. The residuals analysis strongly indicates a more complex demographic history for these populations—such as "phantom" admixture, population-specific bottlenecks, or other intricate dynamics—that is not adequately explained by a simple three-ancestor model.

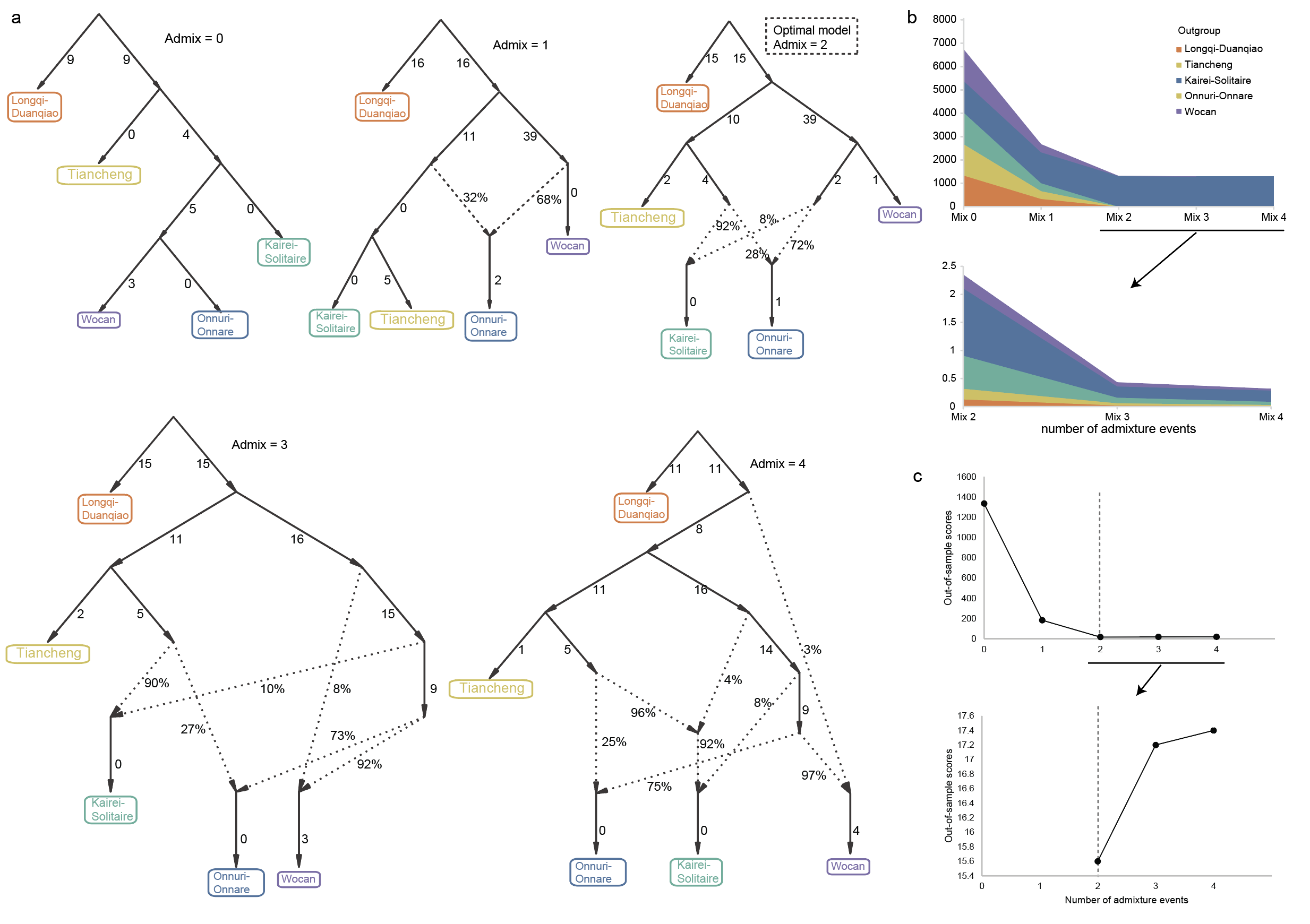

**Figure S5. Population history inference of the scaly-foot snail, *Chrysomallon squamiferum*.** **a** Population history of *C. squamiferum* inferred using qpGraph. Solid lines indicate genetic drift, with lengths proportional to the amount of genetic differentiation (*F*_ST_). Dashed lines represent admixture events, with percentages indicating the inferred proportion of gene flow. We marked the optimal model (Admix = 2) in the figure. **b** Model fit scores for different outgroup populations. The Score indicates the goodness of fit between the model and the data, with scores closer to zero signifying a better fit. **c** Model validation and optimization for the number of admixture events. The model was trained on a random 50% of SNPs and tested on the remaining 50%. This cross-validation was repeated for models with different numbers of admixture events. The results show that a model with two admixture events provides the best fit without overfitting.

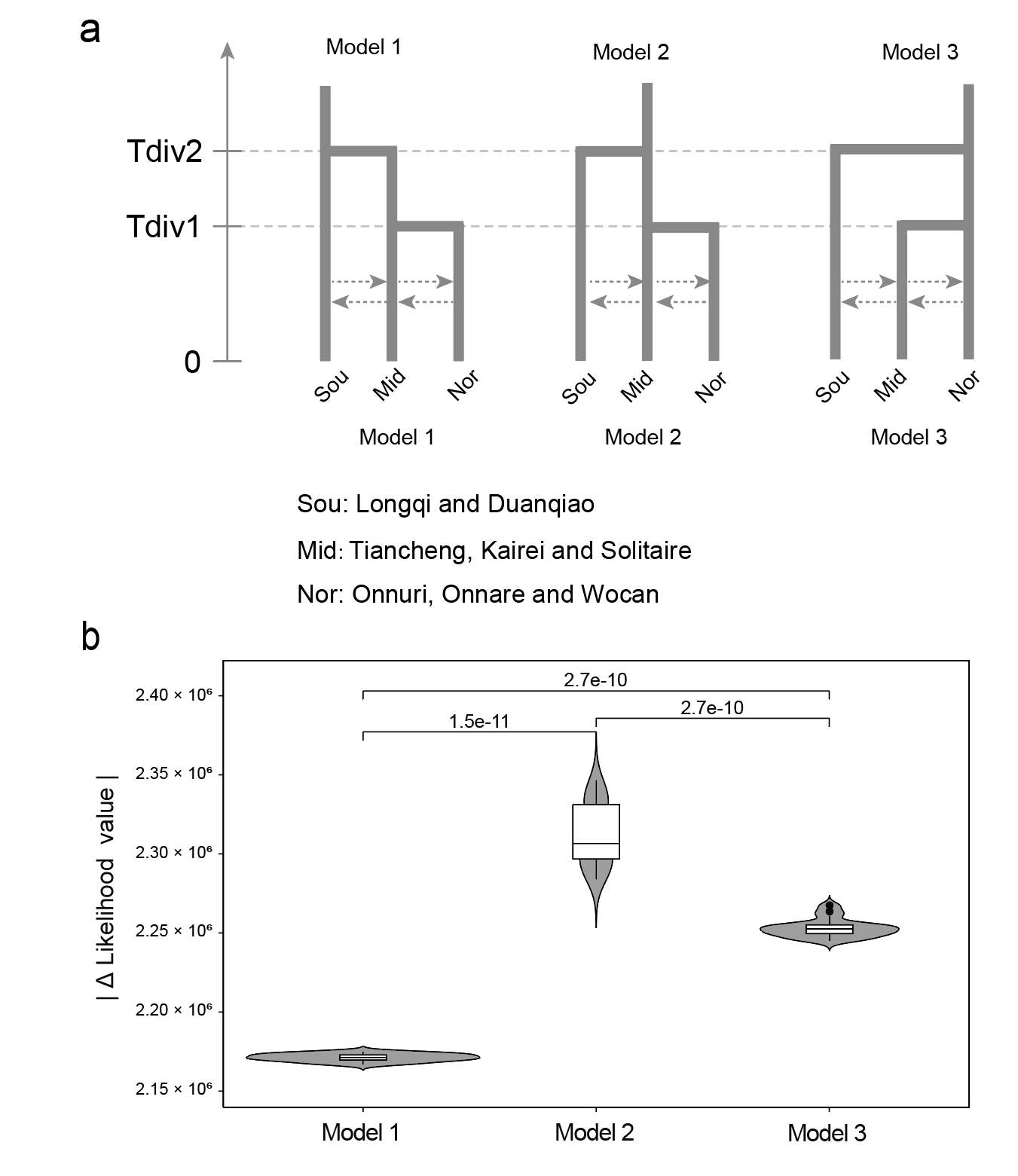

**Figure S6. Demographic model selection for the scaly-foot snail (*Chrysomallon squamiferum*) using Fastsimcoal2**. **a** Schematic representation of three competing demographic models. The models differ in the choice of the basal population (outgroup). The populations are grouped as: 'Sou' (Longqi and Duanqiao), 'Mid' (Tiancheng, Kairei, and Solitaire), and 'Nor' (Onnuri, Onnare, and Wocan). In these models, Tdiv1 is the divergence time between the 'Nor' and 'Mid' groups, and Tdiv2 is the divergence time between the 'Mid' and 'Sou' groups, with the constraint that Tdiv2 ≥ Tdiv1. **b** Model comparison based on delta likelihood scores. The delta likelihood score, where a lower value indicates a better model fit, is plotted for each scenario. The model with the 'Sou' populations as the outgroup was identified as the best fit. The associated *P*-value (<<0.01) confirms that the fit of this model is significantly better than the alternatives.

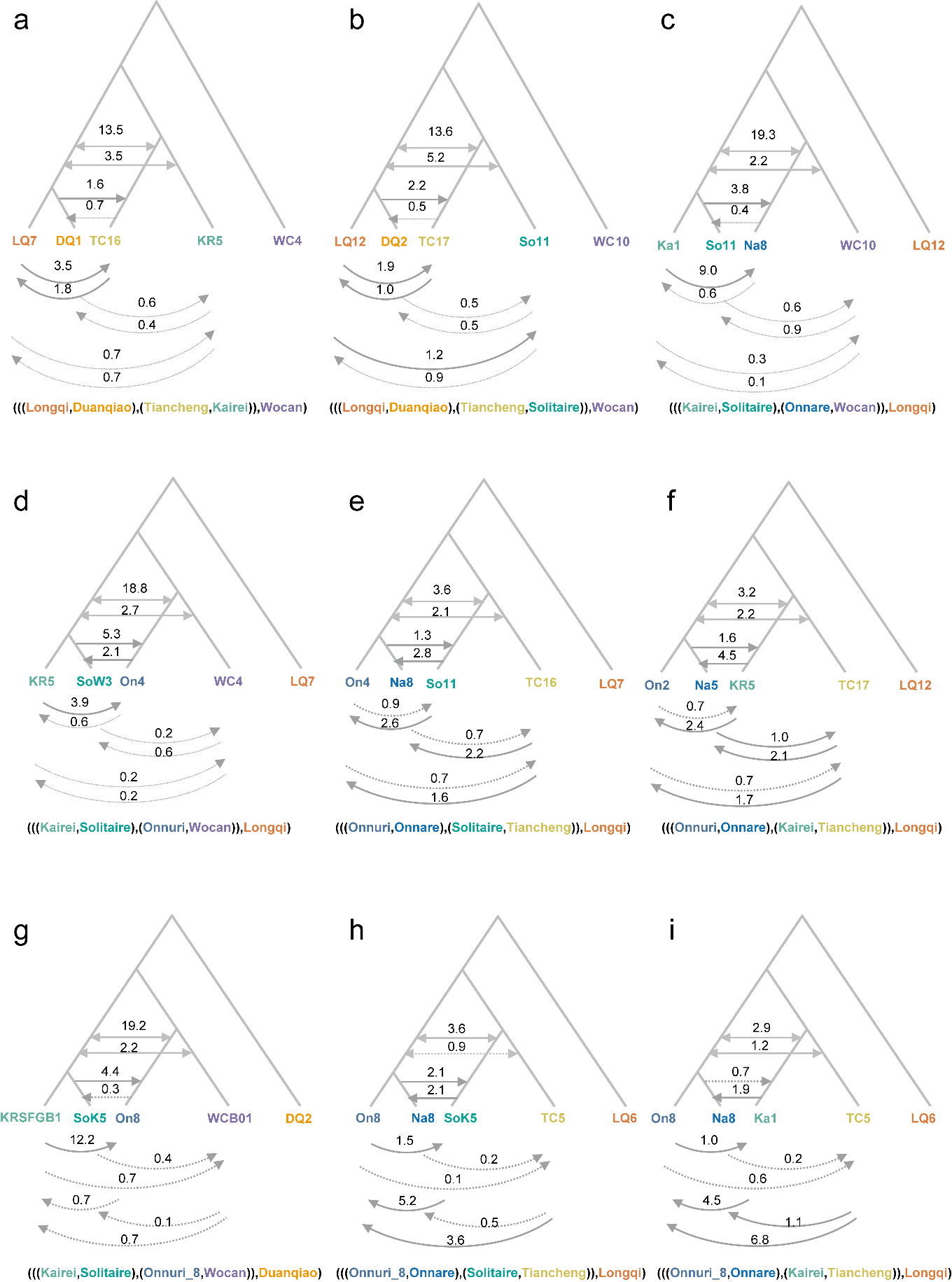

**Figure S7. *Dfoil* test of gene flow direction among scaly-foot snail populations.** The analysis was based on 2024 genomic regions generated from a 200 kb sliding window scan of the genome. Single-headed arrows indicate the inferred direction of gene flow, while double-headed arrows represent an undetermined direction. The numbers indicate the percentage (%) of genomic regions that support the corresponding gene flow direction. Dashed lines signify that gene flow between the two populations is supported by less than 1% of the regions. The individual Onnuri_8 (On8), which was identified as an outlier in the population of Onnuri in the former PCA and Structure analyses, was also included in the *Dfoil* tests in panels **g**, **h**, and **i** to ensure the robustness of the conclusions.

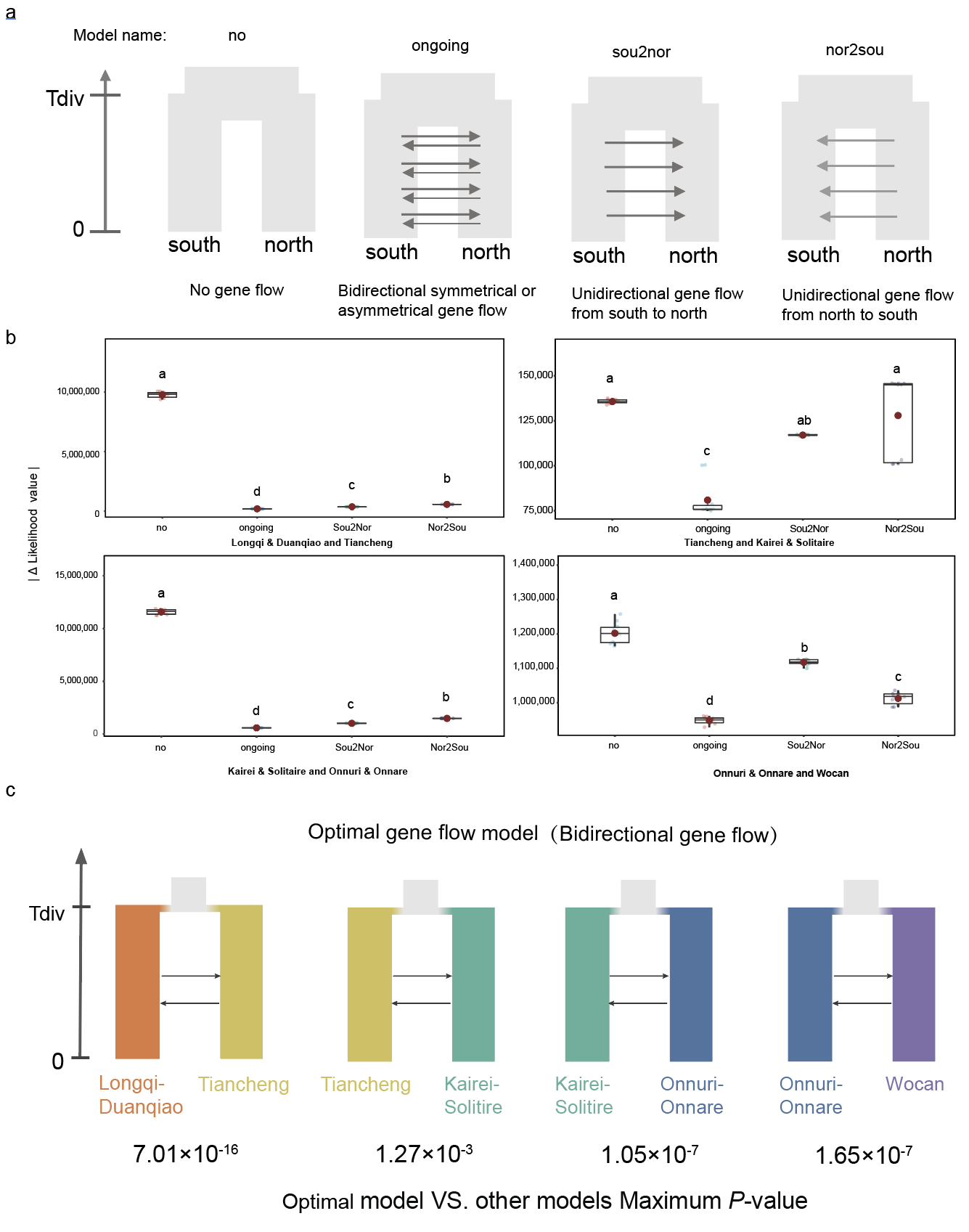

**Figure S8. Gene flow model selection for the scaly-foot snail (*Chrysomallon squamiferum*) using Fastsimcoal2**. **a** Schematic diagrams of the 4 gene flow models tested. The models are categorized based on the directionality of gene flow: "no gene flow" (no gene flow occurs following the population split), "ongoing gene flow" (continuous bidirectional, symmetric or asymmetric, gene flow since population divergence), and "Sou2Nor" and "Nor2Sou" gene flow (unidirectional gene flow since population divergence). **b** The four gene flow models depicted in panel **a** were tested for each pair of the two most geographically proximate population pairs, resulting in a total of four population combinations. The delta Likelihood values were calculated, with the model having the lowest delta Likelihood value considered the best-fit model. Significance is indicated by letters (a, b, c, d) following multiple comparison tests. The results indicate that the "ongoing gene flow" model is the best-fit model for gene flow between the four populations. **c** Gene flow model selection for the scaly-foot snail using Fastsimcoal2. Arrows indicate the direction of gene flow. The results indicate that the continuous bidirectional gene flow model is the best-fit model (*P*-value < 0.01) for gene flow between the four groups.

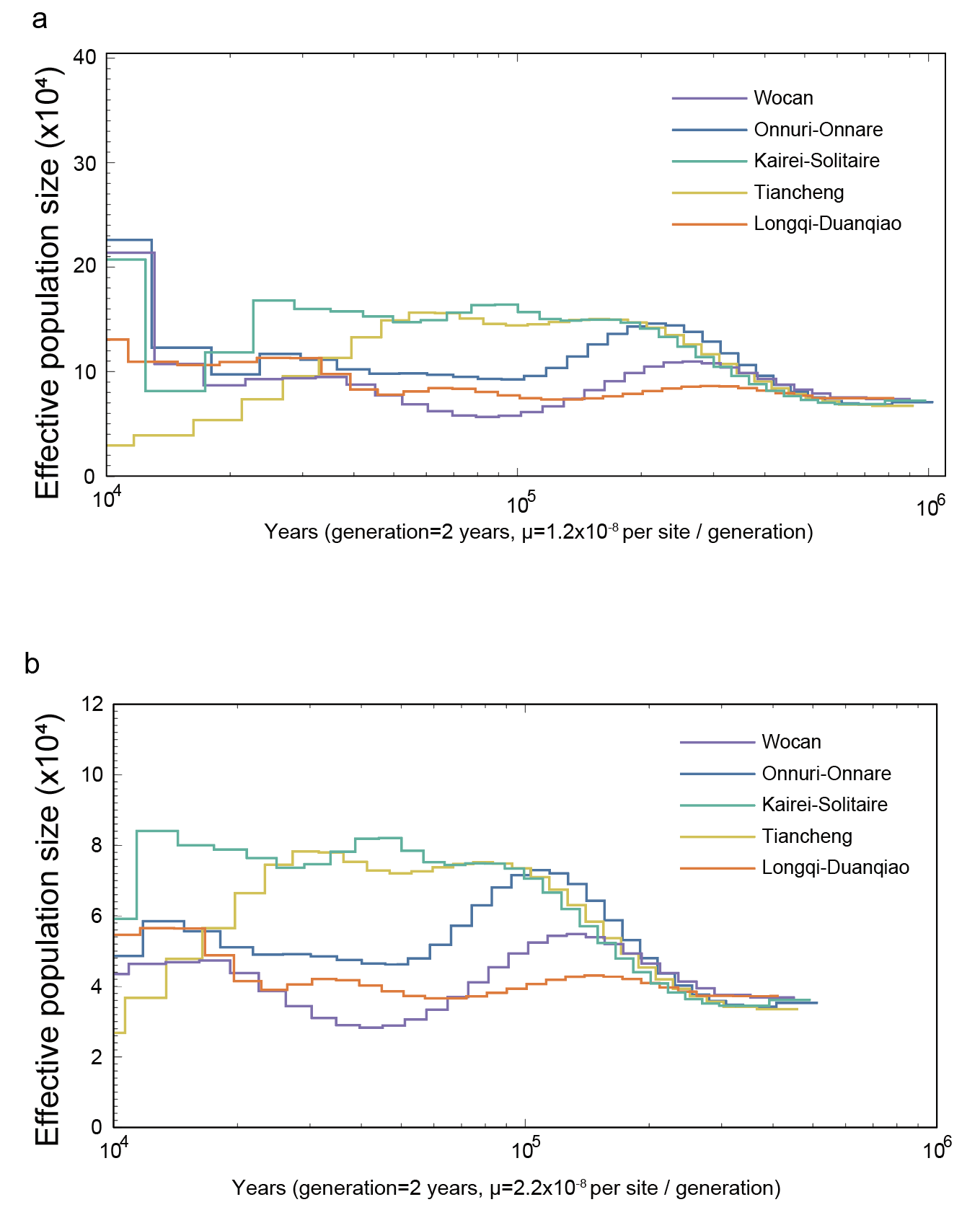

**Figure S9. Historical effective population size (*N*ₑ) of the scaly-foot snail (*Chrysomallon squamiferum*) inferred by PSMC analysis.** The analysis was performed on the individual with the highest sequencing depth from each genetic group. A generation time of 2 years was assumed. To establish a plausible range for the divergence times among the scaly-foot snail populations, the analysis was repeated using four different mutation rate (μ): **a** μ = 1.2×10^−8^, **b** μ = 2.2×10^−8^ per site per generation. The results show that while the absolute scales of Nₑ and time are dependent on the assumed mutation rate, the overall trajectory of demographic change is consistent across all tested rates.

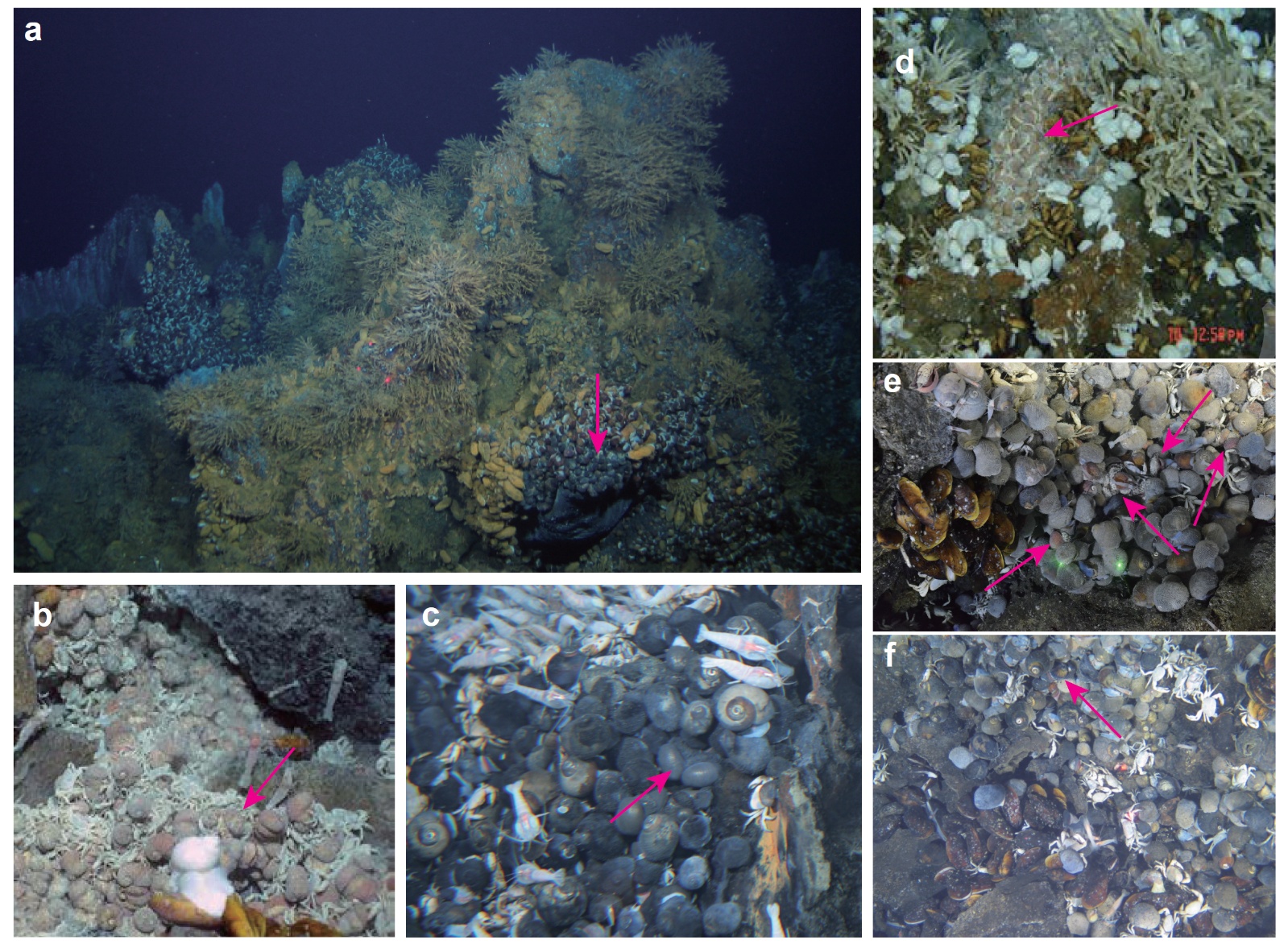

**Figure S10 Habitats of the scaly-foot snails in various hydrothermal vent fields in the Indian Ocean. a** scaly-foot snail population in the Longqi hydrothermal area; **b** Tiancheng hydrothermal area; **c** Kairei hydrothermal area; **d** Solitaire hydrothermal area; **e** Onnare hydrothermal area; **f** Wocan hydrothermal habitat, where only one Scaly-foot Snail can be identified in the photo. The scaly-foot snails are indicated by the red arrows in the figure.
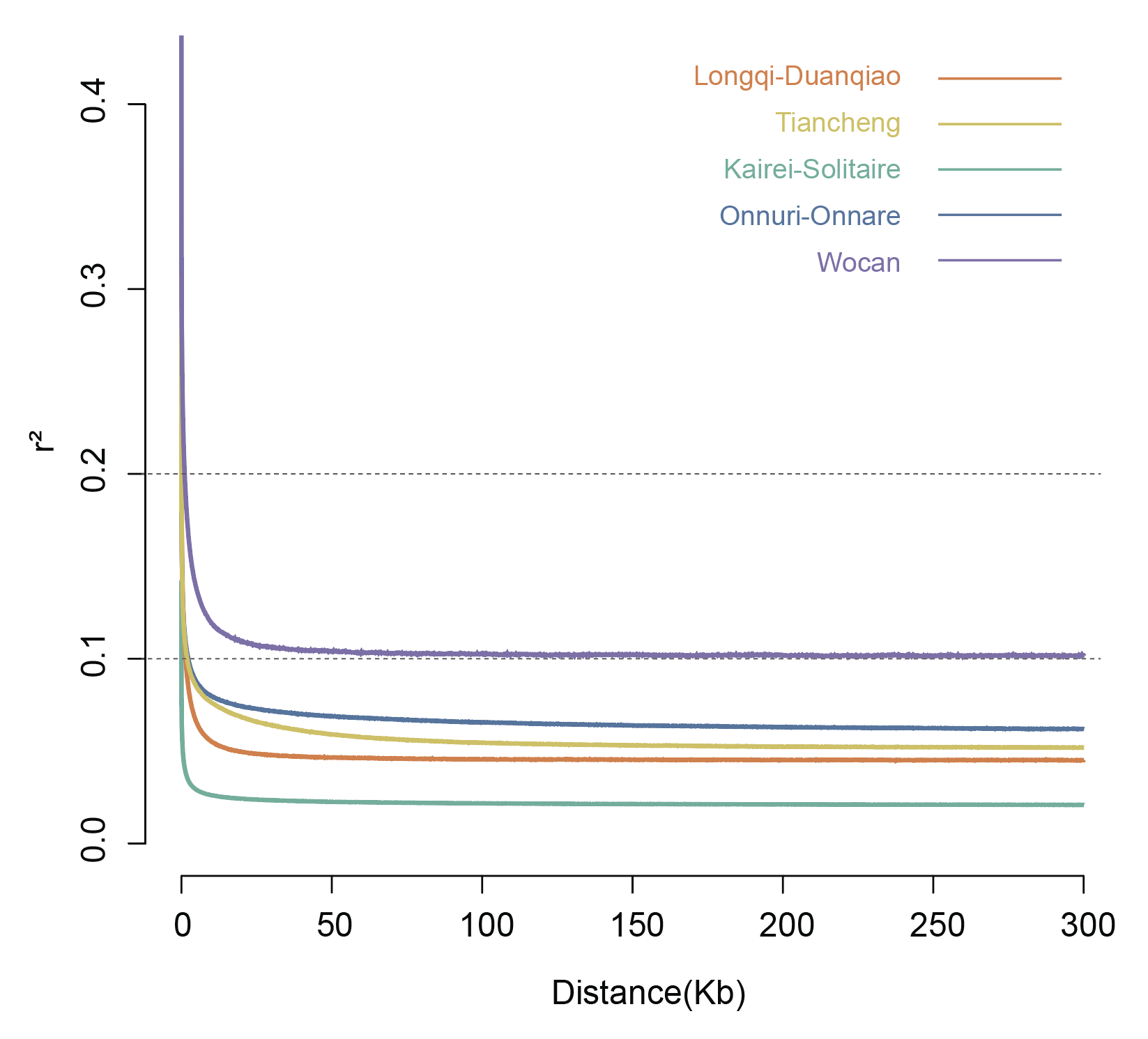

**Figure S11 Figure 1. Decay of linkage disequilibrium (LD) in scaly-foot snail populations.** The average *r*^2^ value is plotted against physical distance (Kb). The plot compares five population groups, showing substantially slower LD decay in the Wocan population compared to the others.

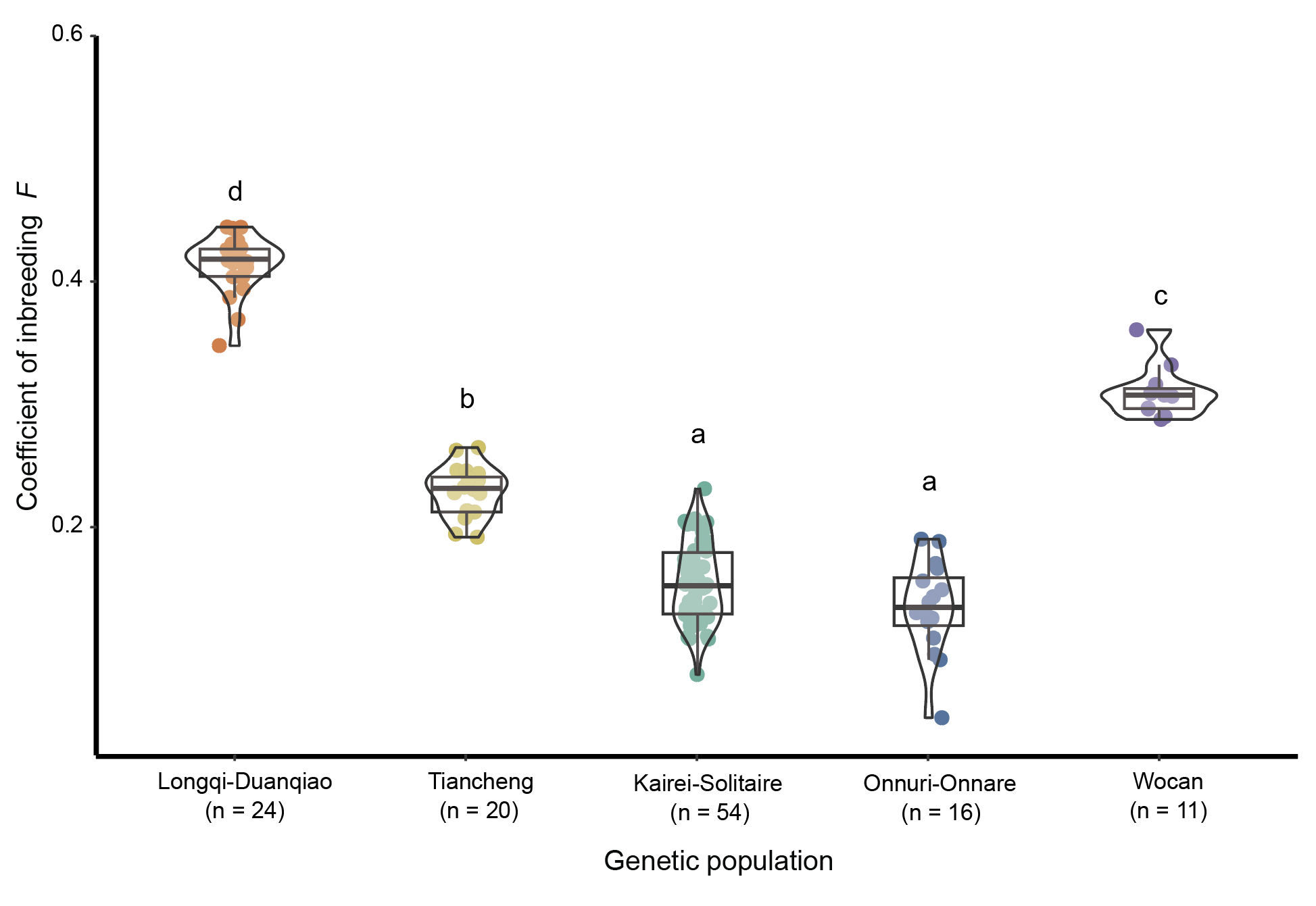

**Figure S12. Inbreeding coefficients of five genetic populations of Scaly-foot Snails.** The results showed that the inbreeding coefficients of Longqi-Duanqiao and Wocan populations were significantly higher than those of the other populations.
